# Countervailing effects of cyclin isoforms A and B during mitosis

**DOI:** 10.64898/2026.01.30.702807

**Authors:** Shrea Bural, Duane A. Compton

**Affiliations:** Department of Biochemistry and Cell Biology, Geisel School of Medicine at Dartmouth Dartmouth Cancer Center, Geisel School of Medicine at Dartmouth, Lebanon, NH

**Author notes:** Lead contact: Department of Biochemistry and Cell Biology, Geisel School of Medicine at Dartmouth, WTRB Level 6, 1 Medical Center Drive, Lebanon, N.H. 03756 USA.

**Keywords:** kinetochore, mitosis, cyclin, cyclin-dependent kinase, kinetochore-microtubule

## Abstract

Faithful chromosome segregation requires the spatial and temporal remodeling of cell structures driven largely by cyclin-dependent kinase (Cdk) activity. In some experimental systems the timely elevation of one cyclin isoform is sufficient to support mitotic entry and progression. In human somatic cells, however, three cyclins – Cyclin A2, Cyclin B1, and Cyclin B2 – are present at mitotic entry, and their distinct contributions during mitosis remain largely unclear. We demonstrate that Cyclin A2 promotes the kinetochore localization of Cyclin B1, Cyclin B2 and the fibrous corona component CENPF, while suppressing recruitment of the kinetochore-microtubule (k-MT) stabilizer Astrin. Extending Cyclin A2 into metaphase by expressing a non-degradable mutant causes persistent kinetochore localization of Cyclin B1, Cyclin B2, and CENPF. Conversely, Cyclin B1 limits kinetochore localization of Cyclin B2 and CENPF, promotes Astrin recruitment, and stabilizes k-MT attachments and Cyclin B1 overexpression further reduces kinetochore localization of CENPF during prometaphase. These findings reveal an interdependence among mitotic cyclins in early mitosis and the countervailing activities of Cyclin A2 versus Cyclin B1/B2 in regulating key mitotic events. We propose that this circuitry enforces a switch-like transition from prometaphase – marked by corona assembly and high k-MT turnover – to metaphase – marked by corona disassembly, stabilization of end-on k-MT attachments, and eventual spindle assembly checkpoint satisfaction – to choreograph the structural changes required to ensure faithful chromosome segregation.

## Introduction

The mechanical separation of replicated chromosomes during mitosis requires dramatic rearrangement of cell contents (Chen *et al*, 2025). These structural rearrangements must be tightly coordinated – both spatially and temporally – to ensure that chromosome segregation occurs with high fidelity to preserve genome integrity in daughter cells (Scholey *et al*, 2003). Cells rely on protein phosphorylation as a primary mechanism to catalyze these structural changes during mitosis because this post-translation modification is rapid and reversible through kinase and phosphatase activities (Cordeiro *et al*, 2018). Cyclin-dependent kinase (Cdk) activity is essential to initiate mitosis in eukaryotes with the timing of Cdk activation depending on association with non-catalytic cyclin subunits (Murray, 1993; Maller, 1991; Nurse, 2002; Pines, 1995). It has been demonstrated under specific experimental conditions that the timely elevation of one cyclin isoform is sufficient to activate Cdk to initiate mitosis and that mitotic exit ensues upon the subsequent ubiquitin-mediated destruction of that cyclin (Fisher & Nurse, 1996). In contrast, human somatic cells enter mitosis with three cyclin isoforms – Cyclin A2, Cyclin B1, and Cyclin B2 – and the distinct contribution that each cyclin isoform makes toward regulating Cdk activity to choreograph structural changes such as kinetochore maturation and k-MT attachment during mitosis remains poorly understood. To date, much of the experimentation to explore the relative contributions of three cyclin isoforms in human cells has focused on the role of Cyclin B1 for regulating mitotic entry, nuclear envelope breakdown and the spindle-assembly checkpoint, the role of Cyclin B2 in the spindle-assembly checkpoint, and the roles of Cyclin A2 on the G2-M transition, nuclear envelope breakdown, and overall mitotic timing/duration (Gong & Ferrell, 2010a; Furuno *et al*, 1999; Gong *et al*, 2007; den Elzen & Pines, 2001, 2; Valles *et al*, 2024; Allan *et al*, 2020; Liu *et al*, 2022; Fung & Poon, 2005; Murray, 2004; Crncec *et al*, 2025; Pines, 1995). There has been less attention to the potential for cyclin isoform-specific roles during mitotic events that are critical for chromosome alignment and faithful segregation. We previously showed that Cyclin A2 is required during prometaphase (early mitosis) to regulate Polo-like kinase 1 activity and to destabilize kinetochore-microtubule (k-MT) attachments to foster correction of erroneously oriented k-MT attachments (Dumitru *et al*, 2017; Kabeche & Compton, 2013). These observations raise the possibility that key structural events occurring during mitosis are subject to cyclin isoform-specific Cdk regulation.

## Results and Discussion

To test the impact of loss of individual cyclin isoforms on specific mitotic events, we utilized siRNA to deplete Cyclin A2, Cyclin B1 or Cyclin B2, the three cyclin isoforms present during mitosis in human somatic cells (Cyclin A1 and Cyclin B3 expression are testes- and embryo-specific, respectively) (Liu *et al*, 2022; Wolgemuth *et al*, 2004). We elected to use siRNA to knockdown expression of each cyclin isoform to examine the effects of their acute loss to avoid compensatory mechanisms that have been described arising through clonal expansion of cells or organisms following genetic ablation of specific cyclin genes (Kalaszczynska *et al*, 2009; Crncec *et al*, 2025). Further, we conducted many experiments in two cell lines (Hela and U2OS) to ensure reproducibility, because non-transformed cells (e.g. RPE1) do not enter mitosis efficiently following loss of cyclin A2 (Hégarat *et al*, 2020). In both HeLa and U2OS cell lines, quantification of cyclin levels in asynchronous cells following siRNA transfection demonstrated significant reduction in expression of the specific cyclin isoform targeted and little to no change in the continued expression of non-targeted cyclin isoforms (Fig S1A&B). Quantification of images of cyclin isoform levels in individual cells that entered mitosis following siRNA transfection also demonstrated significant reduction of the specific cyclin isoform in mitotic cells targeted by the siRNA (Fig S1C). There were significant numbers of cells staining positive for histone H3 Ser10 phosphorylation indicating that these treatments did not prohibit mitotic entry although the mitotic index increased slightly following Cyclin A2 knockdown and decreased slightly following either Cyclin B1 or B2 knockdown which was expected based on previously published work (Fig S1A,B & D) (Crncec *et al*, 2025; Gong & Ferrell, 2010a; Valles *et al*, 2024; Liu *et al*, 2022; Hégarat *et al*, 2020; Chen *et al*, 2008). Additionally, we demonstrate overexpression of mCherry-tagged Cyclin B1 in asynchronous cells using immunoblotting (Fig S2A) and validate the increase in Cyclin B/Cdk1 activity in mitotic cells based on the increase in staining intensity with an antibody specific to the phospho-acceptor site threonine 31 of Hec1 (Fig S2B) (Kucharski *et al*, 2022). We also used immunoblot analysis to verify overexpression of the mCherry-tagged non-degradable ΔN97 mutant version of Cyclin A2 (den Elzen & Pines, 2001) and imaging of single mitotic cells to verify its expression during prometaphase and persistence into metaphase (Fig S2C&D). We also demonstrate mCherry-tagged Cyclin B1 overexpression in U2OS cells that co-express photoactivatable-GFP alpha tubulin (Fig S2C). For clarity throughout, graphical data for control cells are depicted in grey, cells with altered levels of Cyclin A2, Cyclin B1, and Cyclin B2 are depicted in blue, green and magenta, respectively.

We and others previously demonstrated that high mitotic fidelity relies on a stepwise increase in k-MT attachment stability during the transition from prometaphase to metaphase with Cyclin A2 promoting k-MT detachment during prometaphase to facilitate correction of erroneous k-MT attachments (Kabeche & Compton, 2013; Zhang *et al*, 2017). To test if Cyclin B1 or Cyclin B2 have roles in regulating k-MT attachment stability we quantified microtubule turnover rates in live cells using fluorescence dissipation after photoactivation in U2OS cells stably expressing photoactivatable GFP-tagged alpha-tubulin (Fig 1A-C). Differential interference contrast optics was used to define mitotic cells in prometaphase or metaphase based on chromosome alignment (Fig 1B) and dissipation of fluorescence intensity fit a double exponential curve (R^2^>0.99) with the unstable population representing non-k-MT and the more stable population representing k-MT (Zhai *et al*, 1995) (Fig 1A). The stability of k-MTs, as expressed by half-life of the change in fluorescence intensity of the marked region, in control cells is significantly lower in prometaphase compared to metaphase as expected (Kabeche & Compton, 2013) (Fig 1C). Cells depleted of either Cyclin B1 or Cyclin B2 demonstrated no significant difference in k-MT attachment stability during prometaphase compared to control cells, but a significant reduction in k-MT attachment stability during metaphase compared to control cells indicating that their presence promotes k-MT attachment stability during metaphase. Previously, we demonstrated that k-MT attachments in metaphase cells were destabilized by expression of a non-degradable mutant version of Cyclin A2 that persists into metaphase (Kabeche & Compton, 2013). Here, we observe no significant change to k-MT attachment stability during prometaphase upon overexpression of Cyclin B1 compared to control cells, but reduced k-MT stability in metaphase compared to control cells. This outcome is likely due to the increase in Hec1 phosphorylation at phospho-acceptor site T31 that we observed following Cyclin B1 overexpression (Fig S2B) and the previous observation that excessive Cdk1 activity increases microtubule dynamics to increase chromosome segregation defects (Schmidt *et al*, 2021). We did not observe a significant change in the proportion of microtubules in the unstable and stable populations under any of these experimental conditions. These data demonstrate a cyclin isoform-specific effect on k-MT attachment stability during mitosis with Cyclin A2 destabilizing attachments during prometaphase and Cyclin B1 and B2 stabilizing attachments during metaphase.

**Figure 1:**
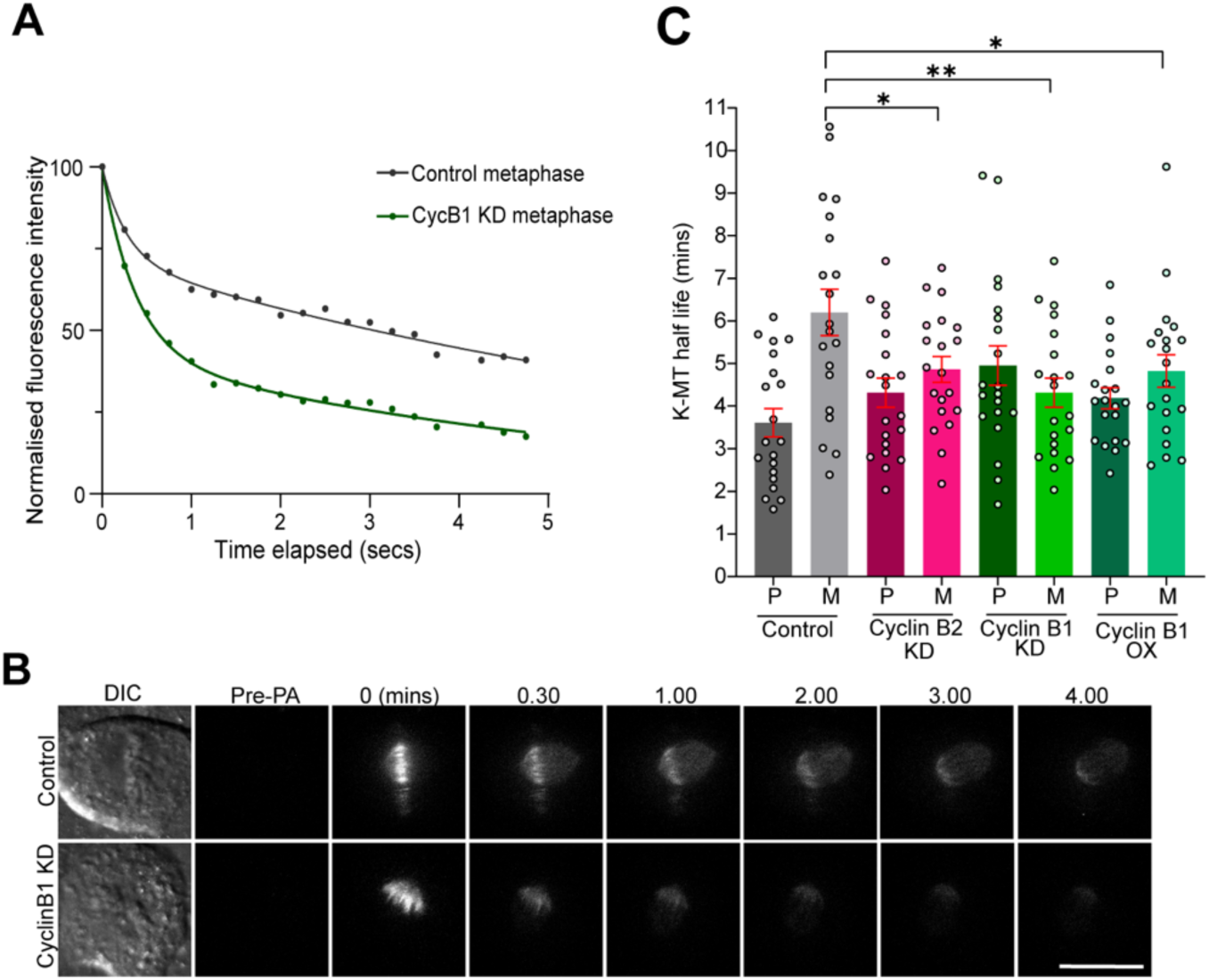
Cyclin isoform-specific regulation of kinetochore-microtubule attachment stability. A. Examples of normalized fluorescence dissipation after photoactivation in metaphase for individual control (grey) and Cyclin B1-depleted (Cyclin B1 KD) (green) cells that are representative of the average k-MT half-life in panel C. B. Representative DIC and fluorescence images (maximum intensity projections) of photoactivated, metaphase control and Cyclin B1-depleted cells. Scale bar, 10 μm. pre-PA, pre-Photoactivation. C. Average k-MT half-life was measured for Control (grey), Cyclin A2-depleted (CyclinA2 KD) (blue), Cyclin B2-depleted (Cyclin B2 KD) (magenta), Cyclin B1-depleted (CyclinB1 KD) (green) and Cyclin B1-overexpressed (CyclinB1 OX) (dark green) cells in both prometaphase (P) and metaphase (M) in U2OS cells expressing photoactivatable GFP-α-tubulin. Error bars indicate SEM; n ≥ 20 cells per condition; One way Anova was performed (\**p* = 0.04 (Control metaphase vs Cyclin B2 KD metaphase), \*\**p* = 0.0017 (Control metaphase vs Cyclin B1 KD metaphase), \**p* = 0.0335 (Control metaphase vs Cyclin B1 OX metaphase).

Astrin conspicuously localizes to kinetochores of chromosomes with bi-oriented spindle attachments (Mack & Compton, 2001; Conti *et al*, 2019; Kern *et al*, 2017; Schmidt *et al*, 2010; Friese *et al*, 2016) and contributes to stabilizing k-MTs during metaphase (Manning *et al*, 2010; Song *et al*, 2021; Geraghty *et al*, 2021; Rosas-Salvans *et al*, 2022). Astrin is a substrate for Cdk phosphorylation (Song *et al*, 2021; Chung *et al*, 2016) and we tested the role of Cyclin A2, Cyclin B1 and Cylin B2 in the temporal control of its kinetochore localization (Fig 2A&B). As expected, Astrin localization at kinetochores increases as control cells transit from prometaphase to metaphase. Cells depleted of Cyclin A2 display a significant increase of Astrin localization to kinetochores during prometaphase compared to control cells and no observable difference in localization during metaphase indicating that Cyclin A2 suppresses Astrin localization during early mitosis. In contrast, cells depleted of Cyclin B1 display a significant decrease in Astrin localization to kinetochores during both prometaphase and metaphase indicating that Cyclin B1 promotes Astrin localization. Cells depleted of Cyclin B2 also displayed a reduction in Astrin localization although the effect was limited to metaphase and the effect size was smaller compared to cells depleted of Cyclin B1. Thus, the temporal control of Astin localization to kinetochores experiences cyclin isoform-specific effects with Cyclin A2 suppressing localization and Cyclin B1 and Cyclin B2 promoting localization.

**Figure 2:**
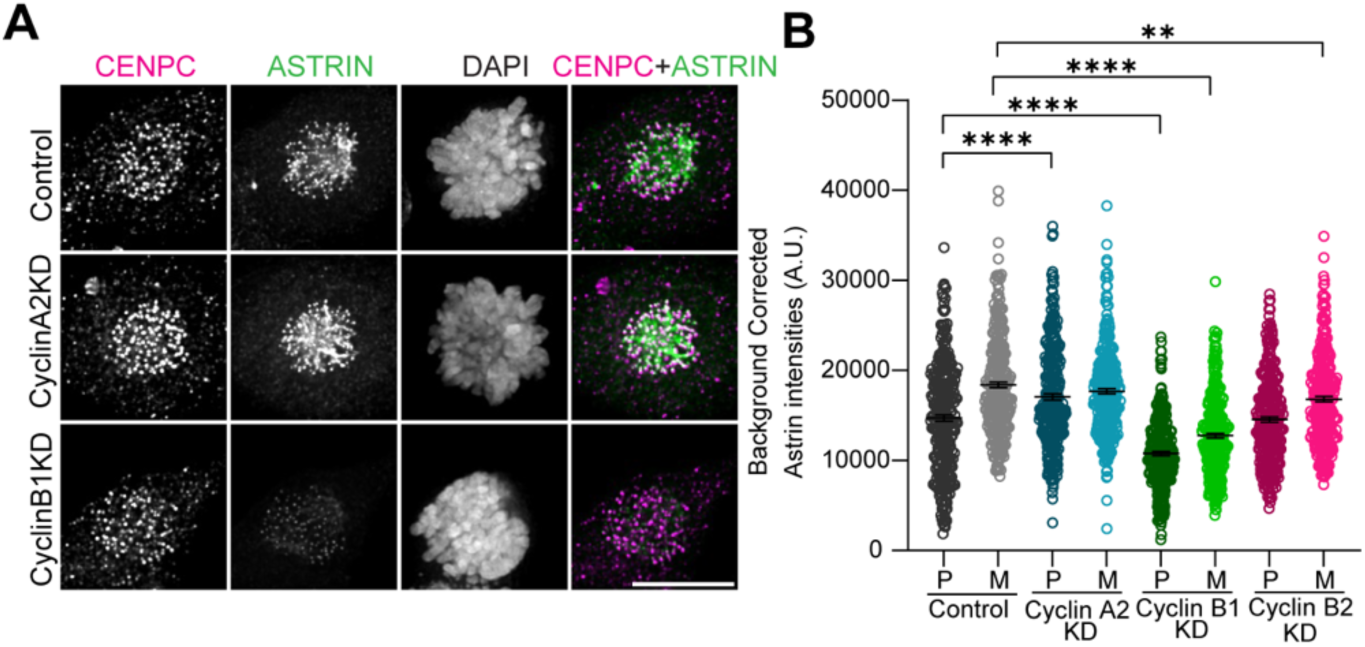
Cyclin isoform-specific regulation of localization of Astrin. A. Immunofluorescence images of Control, Cyclin A2 KD, and Cyclin B1 KD HeLa cells stained for centromere protein C (CENPC; magenta), Astrin (green) and DNA (grey). Scale bar, 10 μm. B. Quantification of Astrin intensities localized at the kinetochores in Control (grey), Cyclin A2 KD (blue), Cyclin B1 KD (green) and Cyclin B2 KD (magenta) cells in both prometaphase and metaphase (N=3, n=300 kinetochores from 30 cells). One-way Anova was performed. \*\*\*\**p* ≤ 0.0001, \*\**p* = 0.0038 using a one-way ANOVA and Šídák’s multiple comparisons test; error bars (black) indicate SEM.

Cyclin A2 is cytosolic throughout the S/G2 transition (Silva Cascales *et al*, 2021) whereas Cylin B1 and Cyclin B2 localize to unattached kinetochores and spindle microtubules (Allan *et al*, 2020; Liu *et al*, 2022; Chen *et al*, 2008; Bentley *et al*, 2007; Alfonso-Pérez *et al*, 2019) during mitosis, primarily during prometaphase. The localization of Cyclin B1 to kinetochores has been shown to be dependent on Cyclin A2 (Kucharski *et al*, 2022), and here we explore additional relationships among cyclin isoforms for kinetochore localization (Fig 3). Cells in prometaphase depleted of Cyclin A2 display a significant reduction in kinetochore localization of Cyclin B1 as expected. Cyclin A2-depleted cells also show a significant reduction in kinetochore localization of Cyclin B2 (Fig 3A&B) indicating that Cyclin A2 promotes localization of both B-type cyclins. Cells in prometaphase depleted of Cyclin B1 show a significant increase in kinetochore localization of Cyclin B2 (Fig 3C) and cells in prometaphase depleted of Cyclin B2 show no observable change in the localization of Cyclin B1 (Fig 3D). This relationship of cyclin-isoform dependent kinetochore localization was replicated in U2OS cells (Fig S3). We reasoned that the timing of Cyclin B1 and Cyclin B2 departure from kinetochores should be altered by preventing the timely destruction of Cyclin A2 during prometaphase. To test this, we expressed mCherry-tagged non-degradable ΔN97 mutant of Cyclin A2 and examined the localization of Cyclin B1 and Cyclin B2 during metaphase (Fig 3E&F). The expression of ΔN97 Cyclin A2 significantly elevated Cyclin B1 and Cyclin B2 localization to kinetochores in prometaphase cells and induced a more persistent localization of both Cyclin B isoforms at kinetochores in metaphase cells. Thus, Cyclin B1 and Cyclin B2 rely on Cyclin A2 presence to promote their localization to kinetochores in early mitosis.

**Figure 3:**
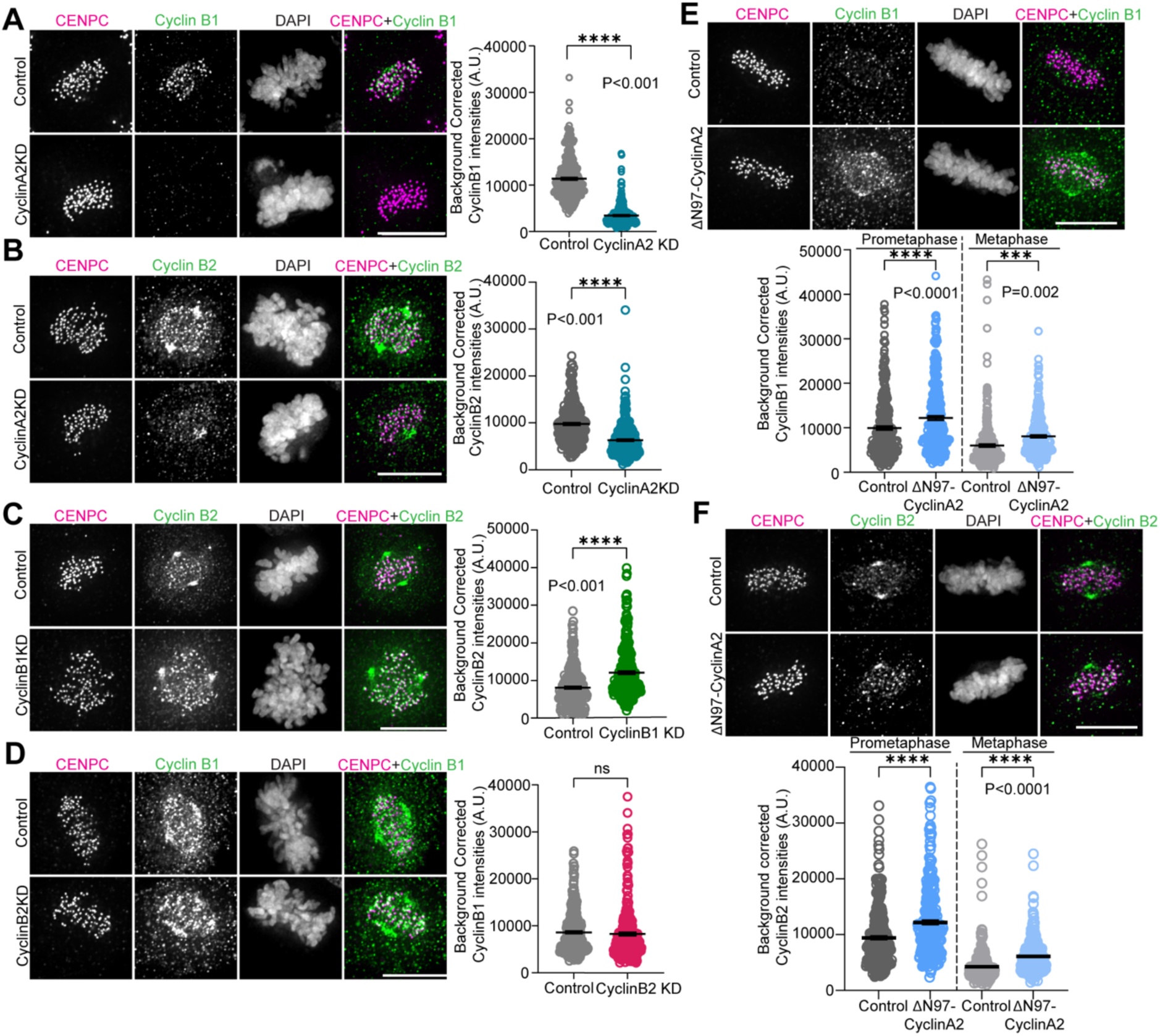
Cyclin isoform-specific regulation of cyclin localization. A. Immunofluorescence images of Control and Cyclin A2 KD HeLa cells stained for CENPC, Cyclin B1 and DAPI (DNA). Quantification of Cyclin B1 intensities localized at the kinetochores in Control (grey) and Cyclin A2 KD (blue) cells (N=3, n=300 kinetochores from 30 cells). Two tailed t-test was performed. B. Immunofluorescence images of Control and Cyclin A2 KD cells stained for CENPC, Cyclin B2 and DAPI (DNA). Quantification of Cyclin B2 intensities localized at the kinetochores in Control (grey) and Cyclin A2 KD (blue) cells (N=3, n=300 kinetochores from 30 cells). Two tailed t-test was performed. C. Immunofluorescence images of Control and Cyclin B1 KD cells stained for CENPC, Cyclin B2 and DAPI (DNA). Quantification of Cyclin B2 intensities localized at the kinetochores in Control (grey) and Cyclin B1 KD (green) cells (N=3, n=300 kinetochores from 30 cells). Two tailed t-test was performed. D. Immunofluorescence images of Control & Cyclin B1 KD stained for CENPC, CyclinB1 and DAPI (DNA). Quantification of Cyclin B1 intensities at kinetochores in Control (grey) and Cyclin B2 KD (magenta) cells (N=3, n=300 kinetochores from 30 cells). Two tailed t-test was performed. E. Immunofluorescence images of Control and nondegradable Cyclin A2 (ΔN97-CyclinA2) expressing cells during metaphase stained for CENPC, Cyclin B1 and DAPI (DNA). Quantification of Cyclin B1 intensities localized at the kinetochores in control (grey) and ΔN97-Cyclin A2 (blue) cells were made in both prometaphase and metaphase (N=3, n=300 kinetochores from 30 cells). One-way anova was performed. F. Immunofluorescence images of Control and nondegradable Cyclin A2 (ΔN97-CyclinA2) expressing cells during metaphase stained for CENPC, Cyclin B2 and DAPI (DNA). Quantification of Cyclin B2 intensities localized at the kinetochores in control (grey) and ΔN97-Cyclin A2 (blue) cells were made in both prometaphase and metaphase (N=3, n=300 kinetochores from 30 cells). One-way Anova was performed. Scale bar, 10 μm for all image panels.

The assembly of the multi-component fibrous corona of the kinetochore is also temporally controlled during mitosis (Cheeseman, 2014; Kops & Gassmann, 2020; Sacristan *et al*, 2018; Wu *et al*, 2023). It is assembled in prometaphase to aid the efficiency of capture of spindle microtubules and is disassembled in metaphase as stable end-on k-MT attachments are established (Chen *et al*, 2025; Magidson *et al*, 2015). Centromere protein F (CENPF) is a key component of fibrous corona that stabilizes microtubule attachments and limits the minus end-directed motor activity of cytoplasmic dynein to strip fibrous corona components from kinetochores (Auckland *et al*, 2020). The stability of the fibrous corona depends on Cdk activity (Sacristan *et al*, 2018) although it remains unknown if it is subject to cyclin isoform-specific effects. To test this, we quantified the kinetochore localization of CENPF in mitotic cells following the manipulation of specific cyclin levels (Fig 4). Cells depleted of Cyclin A2 display a significant reduction in CENPF localized to kinetochores during prometaphase compared to control cells (Fig 4A). In contrast, cells depleted of Cyclin B1 display a significant increase of CENPF localized to kinetochores during prometaphase compared to control cells (Fig 4C). These cyclin isoform-specific effects on CENPF localization were replicated in U2OS cells (Fig S4A&C). Cells depleted of Cyclin B2 also showed slight yet significant increases in CENPF kinetochore localization (Fig S4E&G). Importantly, these cyclin isoform-specific effects on CENPF localization were observed in cells that entered mitosis in the presence of nocodazole indicating that the localization changes are not dependent on microtubule attachments and/or the minus end-directed motor activity of cytoplasmic dynein (Fig 4B&D, Fig S4B&D&F&H). These data posit that cyclin overexpression would inversely affect the quantities of CENPF localized to kinetochores. To test this idea, we quantified CENPF kinetochore localization in cells overexpressing mCherry-tagged Cyclin B1 and observed a significant reduction of CENPF at kinetochores during prometaphase and metaphase (Fig 4E). Further, cells expressing non-degradable mutant ΔN97 Cyclin A2 displayed significant increases in CENPF localization at kinetochores in prometaphase including persistence during metaphase, a stage of mitosis when the fibrous corona is typically disassembled. These observations demonstrate that localization of CENPF to the fibrous corona of kinetochores is subject to cyclin isoform-specific effects whereby Cyclin A2 promotes localization and Cyclin B1 (and to a limited extent, Cyclin B2) suppresses localization.

**Figure 4:**
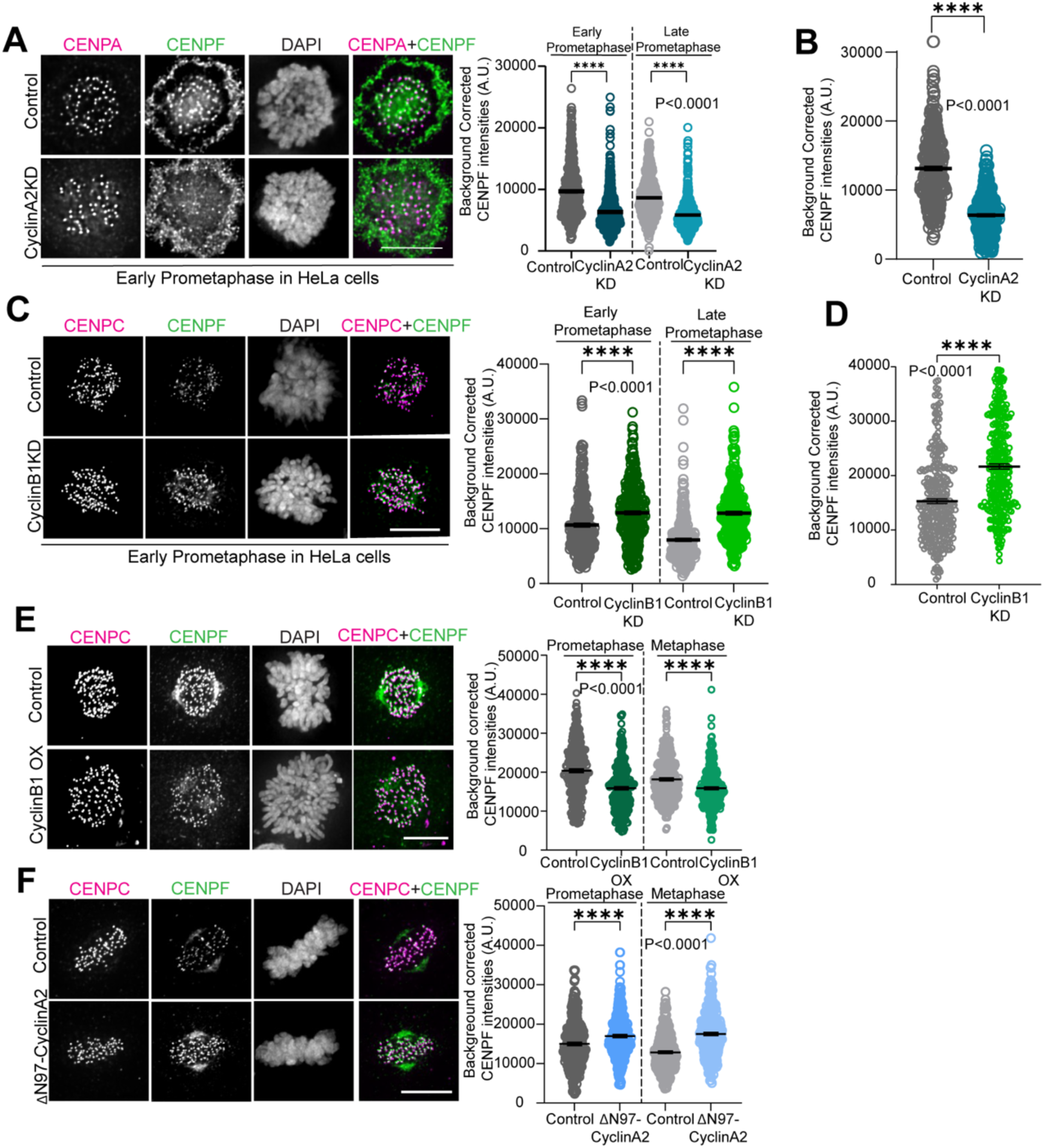
Cyclin isoform-specific localization of CENPF. A. Immunofluorescence images of Control and Cyclin A2 KD cells during prometaphase stained for CENPA, CENPF and DAPI (DNA). Quantification of CENPF intensities localized at the kinetochores in control (grey) and Cyclin A2 KD (blue) cells in early and late prometaphase (N=3, n=300 kinetochores from 30 cells). One-way Anova was performed. B. Quantification of CENPF intensities localized at the kinetochores in control (grey) and Cyclin A2 KD (blue) cells (N=3, n=300 kinetochores from 30 cells). Cells were treated with 3.3μM nocodazole for 2 hours prior to fixation to deplete spindle microtubules. Student t-test was performed. C. Immunofluorescence images of Control and Cyclin B1 KD cells during prometaphase stained for CENPC, CENPF and DAPI (DNA). Quantification of CENPF intensities localized at the kinetochores in control (grey) and Cyclin B1 KD (green) cells in early and late prometaphase (N=3, n=300 kinetochores from 30 cells). One-way Anova was performed. D. Quantification of CENPF intensities localized at the kinetochores in control (grey) and Cyclin B1 KD (green) cells (N=3, n=300 kinetochores from 30 cells). Cells were treated with 3.3μM nocodazole for 2 hours prior to fixation to deplete spindle microtubules. Student t-test was performed. E. Immunofluorescence images of Control and Cyclin B1 overexpressed (OX) cells during prometaphase stained for CENPC, CENPF and DAPI (DNA). Quantification of CENPF intensities localized at the kinetochores in control (grey) and CyclinB1 OX (green) cells in prometaphase and metaphase (N=3, n=300 kinetochores from 30 cells). One-way anova was performed. F. Immunofluorescence images of Control and ΔN97-CyclinA2 expressing cells during metaphase stained for CENPC, CENPF and DAPI (DNA). Quantification of CENPF intensities localized at the kinetochores in control (grey) and ΔN97-Cyclin A2 (blue) cells in prometaphase and metaphase (N=3, n=300 kinetochores from 30 cells). One-way Anova was performed. Scale bar, 10 μm for all image panels.

Taken together, these data show a striking interdependence among cyclin isoforms during early mitosis to choreograph essential structural events with Cyclin A2 promoting the kinetochore localization of both Cyclin B1 and Cyclin B2 (Fig 5). In fostering the kinetochore localization of Cyclin B1 and Cyclin B2, Cyclin A2/Cdk acts as a trigger, or priming kinase, of their function for k-MT attachment error correction via Cyclin B1-dependent phosphorylation of Hec1 (Kucharski *et al*, 2022) as well as their contribution to spindle assembly checkpoint signaling (Allan *et al*, 2020; Liu *et al*, 2022; Gong & Ferrell, 2010b). Further, these data show that all three mitotic cyclin isoforms contribute to the timing of structural changes of the outer kinetochore occurring during mitosis including the localization of the kinetochore fibrous corona component CENPF, stability of k-MT attachments, and the kinetochore localization of Astrin. In this context, all three mitotic cyclin isoforms confer effector kinase function onto Cdk for these mitotic events. However, unlike how Cyclin A2 and Cyclin B1 positively direct Cdk function for nuclear envelope breakdown and chromatin condensation (Gong & Ferrell, 2010b, 2010b; Fung *et al*, 2007), we observe that Cyclin A2 confers countervailing effects on Cdk function compared to Cyclin B1 and Cyclin B2 toward the mitotic events that we examined (Fig 5). This functional opposition likely arises because individual cyclin isoforms (in this case, Cyclin A2 vs Cyclins B1 & B2) direct Cdk activity toward different substrates to affect the same mitotic event in contrasting ways. Although Cyclin A2/Cdk and Cyclin B1(B2)/Cdk may phosphorylate many of the same acceptor sites in mitosis, the idea of cyclin isoform-specific selectivity of phospho-acceptor sites is supported by myriad data including proteomic analysis of protein complexes (Pagliuca *et al*, 2011; Heinzle *et al*, 2025; Al-Rawi *et al*, 2023), structural predictions for Cdk2 substrate selection (Brown *et al*, 2007; Schulman *et al*, 1998; Lowe *et al*, 2002), evidence for Cyclin A2-dependent non-canonical phospho-acceptor site selection (Al-Rawi *et al*, 2023; Pellarin *et al*, 2025), the identification of MYPT1 as a Cyclin A2 isoform-specific substrate in early mitosis (Dumitru *et al*, 2017), and the identification of Cyclin B1-specific substrates in late mitosis (Hégarat *et al*, 2020; Cui *et al*, 2018).

**Figure 5:**
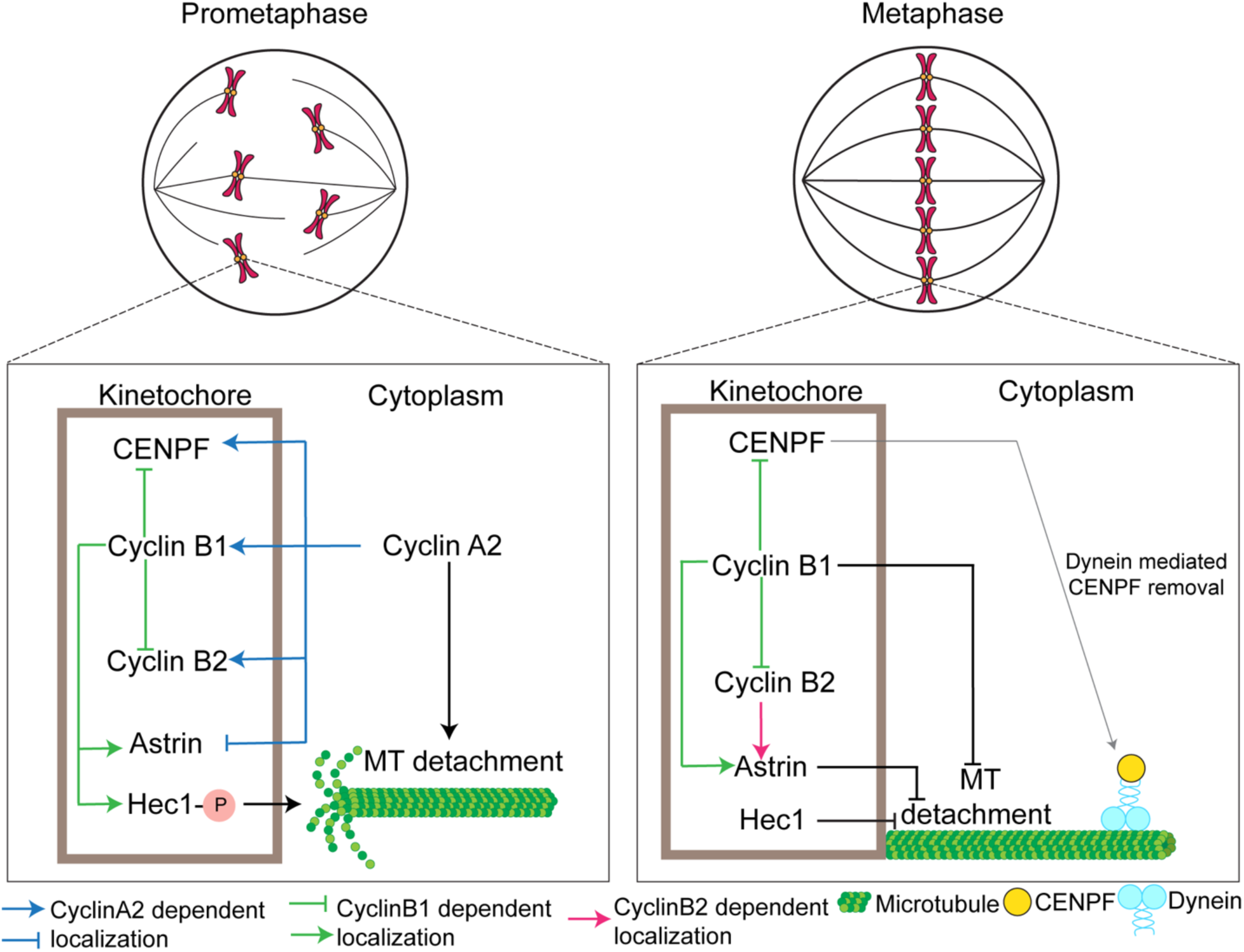
Model for regulation of prometaphase-metaphase transition. As essential co-factors required to stimulate Cdk activity, Cyclin A2 exerts countervailing effects relative to Cyclin B1 and Cyclin B2 on events during prometaphase including k-MT attachment stability, kinetochore localization of Cyclin B2, Astrin and CENPF. Thus, Cyclin A2 aids in k-MT capture and correction of erroneous attachments while simultaneously preparing the stage so that its timely destruction during prometaphase executes a switch-like transition to metaphase.

Finally, evidence shows that both Cyclin-dependent kinases and Aurora B kinase (AurkB) regulate molecular events needed to support faithful chromosome segregation during mitosis including the timely association and subsequent dissociation of centromere/kinetochore components, the correction of k-MT attachment errors (Kucharski *et al*, 2022; Shrestha *et al*, 2017; Broad & DeLuca, 2020; Cimini *et al*, 2006; Liu & Lampson, 2009; Carmena *et al*, 2012; Krenn & Musacchio, 2015) and the spindle assembly checkpoint (D’Angiolella *et al*, 2003; Serpico & Grieco, 2020; Hayward *et al*, 2019; Musacchio & Salmon, 2007; McAinsh & Kops, 2023; Roy *et al*, 2022). Moreover, it’s been demonstrated that Cdk activity can promote centromere localization of Aurora B kinase via Haspin kinase (Kabeche & Compton, 2013; Tsukahara *et al*, 2010) while AurkB activity promotes Cdk kinetochore localization via MPS1 kinase (Valles *et al*, 2024; Cheeseman, 2014; Ali & Stukenberg, 2023) creating a positive feedback loop between the Cdk and AurkB signaling pathways. This molecular circuitry is consistent with a bistable switch (Ferrell, 2002) to regulate the prometaphase to metaphase transition with irreversibility for this specific cell cycle transition introduced when Cyclin A2/Cdk activity drops below a critical threshold due to ongoing ubiquitin-mediated destruction of Cyclin A2 during prometaphase (den Elzen & Pines, 2001). We speculate that the overlap between Cdk and AurkB activities in regulating events at kinetochores during this cell cycle phase transition adds robustness to this circuit because the distributed signaling contributed by Cyclin A2/Cdk activity could dampen potential noise in signaling emanating from the chromosome-autonomous activity of AurkB (den Elzen & Pines, 2001; Gong & Ferrell, 2010b; Pellarin *et al*, 2025; Godek *et al*, 2015; Ali & Stukenberg, 2023).

## Methods

### Cell Culture

HeLa (ATCC, CCL-2) and U2OS (ATCC, HTB-96) cells were grown in DMEM (#15-017-CM, Corning) supplemented with 10% FCS (#82013-586; HyClone), 250 µg/l Amphotericin B (#82026-728; VWR), 50 U/ml penicillin, and 50 µg/ml streptomycin (#15140122; Thermo Fisher Scientific). Cell lines were validated as *Mycoplasma* free (Sigma-Aldrich, Mycoplasma Kit). Cell lines were cultured in a 37°C-humidified environment with 5% CO_2_.

For photoactivation experiments, U2OS cells stably expressing photoactivatable GFP-α-tubulin (Kabeche and Compton, 2013) were cultured in Dulbecco’s Modification of Eagle’s Medium (DMEM, #15-017-CM, Corning) supplemented with 10% FBS (Hyclone), 250 μg/L Amphotericin B (#82026-728; VWR), 50 U/mL penicillin (Mediatech), and 50 μg/mL streptomycin (#15140122; Thermo Fisher Scientific). Cells were grown in incubators at 37°C-humidified environment with 5% CO2.

### Drug treatments

Nocodazole (VWR #80058-500) was used at 3.3 μM for 2 hours.

### Plasmids

Mcherry-Cyclin B1 was obtained from Addgene (Cat# 26063). Mcherry N-97 Cyclin A2 was previously cloned as mentioned in Kabeche et. al, 2012.

### Transfections

For generating knockdown cells, 6-8 x 10^4^ HeLa cells or 4-6 x 10^4^ U2OS cells were plated on 18mm glass coverslips in 12 well plates with DMEM supplemented with 10% FCS, Amphotericin B, pencillin and streptomycin. Cells were transfected 24 hours (HeLa cells) or 36 hours (U2OS cells) after plating with 150 nM Cyclin A2-CCNA2, (Silencer Select Validated; 5′-GA UAUACCCUGGAAAGUCUtt-3′) (Thermo Fisher Scientific, ID: s2513), 100nM ON-TARGETplus Human Cyclin B1-CCNB1 siRNA smartpool (#L-003206-00-0020, Horizondiscovery-Dharmacon) or 100nM ON-TARGETplus Human Cyclin B2-CCNB2 siRNA smartpool (#L-003207-00-0020, Horizondiscovery-Dharmacon) with JetPrime (#89129-922, VWR) following manufacturer’s protocol for siRNA transfection on coverslips in a 12 well plate. Control transfections were performed using JetPrime reagent without any siRNA. 48 hours post transfection; cells were collected for lysate preparation or fixed for Immunofluorescence (IF).

Lipofectamine 3000 (Lipo3000) was used to transfect HeLa cells and U2OS-PA-Tubulin cells with Mcherry-CyclinB1 and Hela cells Δ□N97 -CyclinA2 mcherry. 6-8 x 10^4^ cells were plated on 18mm glass coverslips in 12 well plates with DMEM supplemented with 10% FCS, AmphotericinB, pencillin and streptomycin. 24 hours after plating the cells DMEM was replaced with Opti-MEM (#31985070, Corning) and Lipo3000 along with the plasmid following manufacturer’s protocol for plasmid transfection on coverslips in a 12 well plate. Control transfections were performed using Lipo3000 without any plasmid. Following 4-6 of transfection the Opti-Mem solution was replaced with supplemented DMEM. 48 hours post transfection; lysates were either prepared for Western blot or fixed for IF.

### Antibodies

Primary antibodies for IF are as follows: Rabbit anti-Astrin (Mack and Compton, 2001; 1:1000 IF), mouse anti-CENPC (Abcam, ab50974; 1:100 IF), Guinea pig anti-CENPC (1:1000; MBL; PD030, RRID:AB_10693556), mouse anti-CENP-A (1:500; Thermo Fisher Scientific; MA1-20832, RRID:AB_2078763), mouse anti-Cyclin A2 (1:400; GeneTex; GTX634420, RRID:AB_2888449), rabbit anti-Cyclin B1 (1:200; Cell Signalling; #4138, RRID:AB_2072132), rabbit anti-Cyclin B2 (1:500;abcam; ab185622), Rat anti-Tyrosinated Tubulin (Tubulin; 1:2500; Novus; NB600-506, RRID:AB_10078394), Rat anti-Tyrosinated Tubulin (Tubulin; 1:500; Santa Cruz; sc-53029, RRID:AB_793541), Rabbit anti-CENPF (1:400; Abcam; ab5, RRID:AB_304721) and Mouse anti-Hec1 (1:1,000; Santa Cruz; C-11).The rabbit anti-Hec1 pT31 antibody was generated by Pacific Immunology (Kucharski et. al, 2022; 1:100,000 IF). Secondary antibodies for IF are highly cross-adsorbed Alexa Fluor 488, 594, or 647 antibodies, raised in donkey or goat, against mouse, rabbit, rat or guinea pig (Thermo Fisher Scientific; 1:1000 IF). DAPI was used to stain DNA (500 ng/ml; Sigma-Aldrich; D9542).

Primary antibodies for Western blotting are as follows: Mouse anti-Cyclin A2 (1:2500; Santa Cruz; sc751), Cyclin B1 (1:600; Proteintech; #55004-1-AP), Cyclin B2 (1:1000; abcam; ab185622), GAPDH (1:1000; Santa Cruz Biotechnology; sc-365062), H3pS10 (1:1000; Cell Signaling Technology: #53348). Secondary antibodies for Western blotting are IRDye 680RD Goat anti-Mouse (1:10,000; #926-68070; LI-COR), IRDye 800CW Goat anti-rabbit (1:10,000; #926-32211, LI-COR), and IRDye 800CW Goat anti-mouse (1:10,000; #926-32210; LI-COR).

### Photoactivation

U2OS cells stably expressing photoactivatable GFP-α-tubulin were used to measure k-MT half-life. The cells were imaged with a 100X plan apo, 1.4 NA, oil immersion objective (Nikon) using a QuoromWaveFX-X1 spinning disk confocal system on a Nikon Eclipse Ti microscope equipped with a photometrics evolve delta EMCCD camera, an ILE laser source (Andor Technology), and a Mosaic digital mirror (Andor Technology). Cells were imaged in Fluorobrite DMEM (Thermo Fisher Scientific #A1896701) and placed in a Rose chamber on a heated stage to maintain their temperature at 37℃ during image acquisition. Using differential interference contrast (DIC) microscopy mitotic cells were identified and classified as prometaphase or metaphase based on chromosome alignment. Photoactivation was performed with a 405 nm laser (35% power, 500 ms pulse). A narrow, rectangular region on one half of the spindle was photoactivated, and z-stacks with a range of 6 μm and step size of 1 μm were captured every 15 seconds for 4 minutes. Fiji (ImageJ) was used to quantify the fluorescence dissipation after photoactivation (FDAPA) of maximum intensity projections. Average pixel intensities within the photoactivated region were measured, as well as those in an equally sized area from the nonactivated half of the spindle for background subtraction. To correct for photobleaching, values of fluorescence loss obtained from cells treated with 1 μM Taxol were used. Fluorescence intensities were then normalized to the first time point after photoactivation for each cell. To measure microtubule turnover rates GraphPad software was used to fit the corrected fluorescence intensities at each timepoint to a two-phase exponential decay curve [*F*(*t*)=*A*1*e*−*k*1*t*+*A*2*e*−*k*2*t*], where F(t) is the measured photoactivation fluorescence at time t, A1 is the percentage of fluorescence from the fast decay process representing the non-k-MT population with the decay rate k1, A2 is the percentage of fluorescence from the slow decay process representing the stable k-MT population with the decay rate k2. The turnover half-life for both the slow and fast decay populations of microtubules was then calculated as ln2/k.

### Immunofluorescence

Cells were washed in room temperature DMEM followed by a pre-extraction step to minimize cytoplasmic staining by incubating the cells in 60 mM PIPES, 25 mM HEPES, pH 6.8, 10 mM EGTA, and 2 mM MgCl2 + 1% Triton-X and incubated for 7 mins at 37°C to visualize Astrin, Hec1, Cyclin B1, Cyclin B2, CENPF localized at the kinetochores. The pre-extraction step was skipped for staining cytoplasmic Cyclin A2. Subsequently cells were fixed in 20 mM Pipes, pH 6.8, 10 mM EGTA, 1 mM MgCl2, 0.2% Triton X-100, and 4% formaldehyde for 10 mins at RT. All the steps following fixation was performed at RT. Cells were permeabilized with TBS + 3% BSA+0.5% Triton-X 3×5 mins before blocking in TBS + 3% BSA+0.1% Triton-X for 30 min and incubation with primary antibodies diluted in TBS + 3% BSA+0.1% Triton-X for 2 h at room temperature. Subsequently cells were washed 3×5mins in TBS + 3% BSA+0.1% Triton-X followed by incubation with Alexa-Fluor conjugated secondary antibodies diluted in TBS + 3% BSA+0.1% Triton-X for 1 h at room temperature. Next, cells were washed four times in TBS + 3% BSA+0.1% Triton-X followed by incubation with DAPI (1:2000) diluted in TBS + 3% BSA+0.1% Triton-X. Subsequently cells were washed 3×5 mins in TBS + 3% BSA+0.1% Triton-X followed by 3×5 mins in TBS+0.1% Triton-X. Cells were then mounted in Prolong Gold anti-fade reagent (Invitrogen; Ref: P36934) and imaged.

For Figure 3 (b,c) and Figure S4 (b,d,f,h) cells were treated with 3.33μM Nocodazole and incubated for 2 hours at 37 °C followed by room temperature DMEM washing, pre-extraction, fixation and staining following the protocol as described above.

### SDS-PAGE and Western Blotting

48 hours post-transfections, cells were collected, lysed, and boiled in 4× Laemmli buffer (1M Tris pH6.8, 50% glycerol, 10% SDS, 0.5% bromophenol blue, β-mercaptoethanol) for 10 mins. Proteins were separated on SDS-PAGE gel using stacking gel (4% 29:1 acrylamide: Bis-acrylamide, 125 mM Tris, pH 6.8, 0.1% SDS, 0.1% ammonium persulfate, and 0.1% TEMED and 8–15% 29:1 acrylamide: Bis-acrylamide, 400 mM Tris pH 8.8, 0.1% SDS, 0.1% ammonium persulfate, and 0.1% TEMED) at 80V for 20 mins to stack the proteins followed by 100 V for approximately 2 hours or more until the bromophenol blue has run off, as needed. Proteins were then transferred onto nitrocellulose membranes overnight 30 V under wet conditions in 1× transfer buffer (14.4 g/l glycine, 3.0 g/l Tris, and 20% methanol) (Bio-Rad). The next day, membranes were dried for an hour and rehydrated in 1X TBS for 5 min. Subsequently the membranes were blocked in Intercept (PBS) Blocking Buffer (LICOR; Cat# 927-70001) for 1 h at room temperature (RT). Following blocking, membranes were incubated for 2 hours at RT with primary antibodies diluted in Intercept (PBS) Blocking Buffer + 0.2% Tween (VWR). Next, membranes were washed 4 × with TBS-T (0.1% Tween 20) for 5 min. Membranes were then incubated with secondary antibodies diluted in Intercept (PBS) Blocking Buffer + 0.2% Tween 20 at RT and protected from light. Following incubation with secondary antibodies, membranes were then washed 4 × with TBS-T for 5 min and rinsed with TBS. Membranes were imaged on an Odyssey CLx Digital Imager (LI-COR). Quantification of band signal intensities were performed using Image Studio Lite software (LI-COR). Bar graphs are representative of the protein band intensity relative to GAPDH (loading control).

### Microscopy

All fixed cell immunofluorescence images were acquired using a Nikon Eclipse Ti Inverted microscope using a Plan Apo VC 60x, 1.4 NA, oil immersion objective (Nikon) and equipped with an ORCA-Fusion Gen III sCMOS camera (Hamamatsu) and a SOLA LED light engine (Lumencor). Images were acquired using 0.25 µm steps.

Image deconvolution and contrast enhancement were performed using NIS Batch Deconvolution (Nikon), NIS Elements (Nikon), and ImageJ (NIH). Images shown are maximum intensity projections of selected Z-planes.

### Quantification of Protein expression

For quantification of Astrin, CENPF, Cyclin B1, Cyclin B2, Hec1 and pT31 Hec1 localised at the kinetochores, measurements were taken using the CRaQ Fiji plugin as described in Bodor et al., 2012. Background corrected intensity measurements were calculated as the difference between maximum and minimum intensity values at small areas bounding kinetochores (using the Hec1, CENP-A or CENPC signals to define kinetochore regions) from maximum intensity projections of z-stacks.

Figure images are deconvolved (Nikon Elements) maximum intensity projections from z-stacks of 20–25 imaging planes. An elliptical region of interest (ROI) was drawn to encompass the entire nucleus and then a slightly larger elliptical ROI was drawn to encompass both the nucleus and the adjacent background. The mean background intensity was calculated based on the in-between region of the two ROIs. Expression level of a protein of interest in each nucleus was represented by the background subtracted integrated intensity of the ROI that encompasses the nucleus. Figure panels from different cell lines have different contrast enhancements (Fiji), but contrast within a cell line is consistent for all conditions/mitotic stages, except for the DAPI channel.

### Statistical analyses

All experiments are data from at least three independent biological replicates, unless otherwise specified in the figure legend. Sample sizes are stated in figure legends. All statistical analyses were performed using GraphPad 9. Statistical tests used were Fisher’s Exact test, Two tailed t-test, or one way ANOVA, with multiple comparisons corrections for specified pairs of conditions analyzed. Error bars on graphs represent mean ± SEM. p- values are shown as: ns p > 0.5; *p ≤0.05, **p ≤0.01,***p ≤ 0.001, and ****p ≤ 0.0001 in the figure legends.

### Key resources table

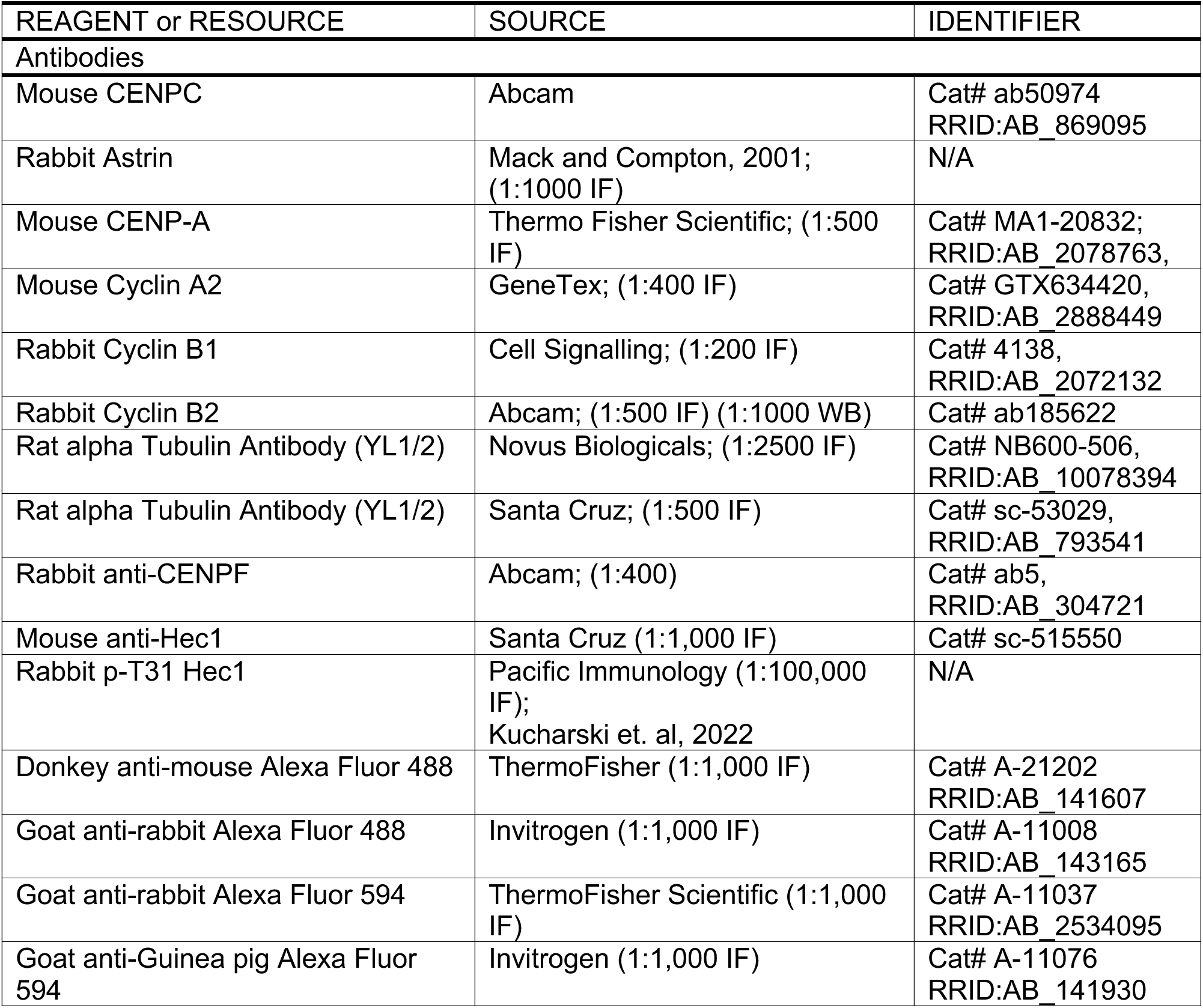
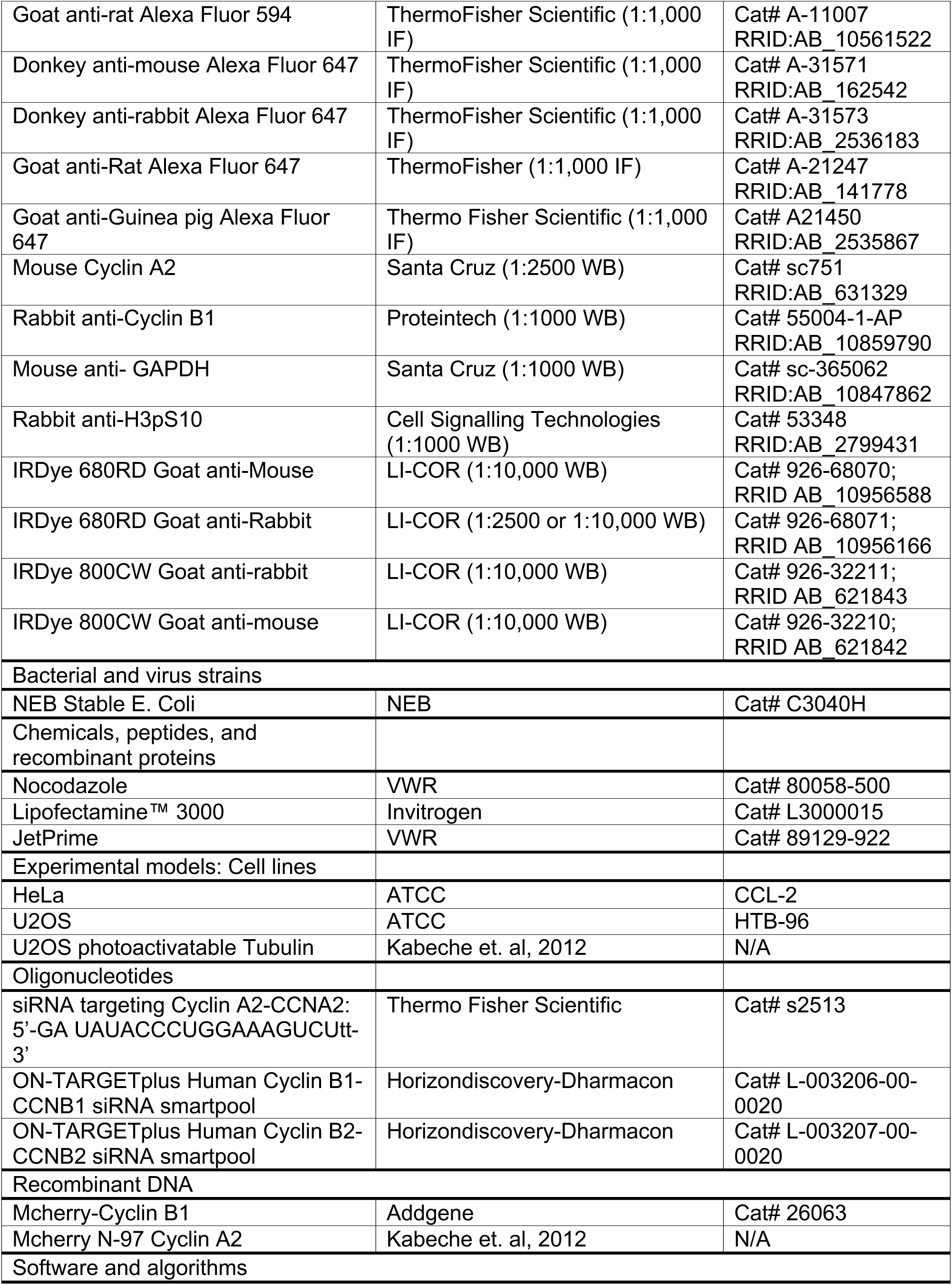
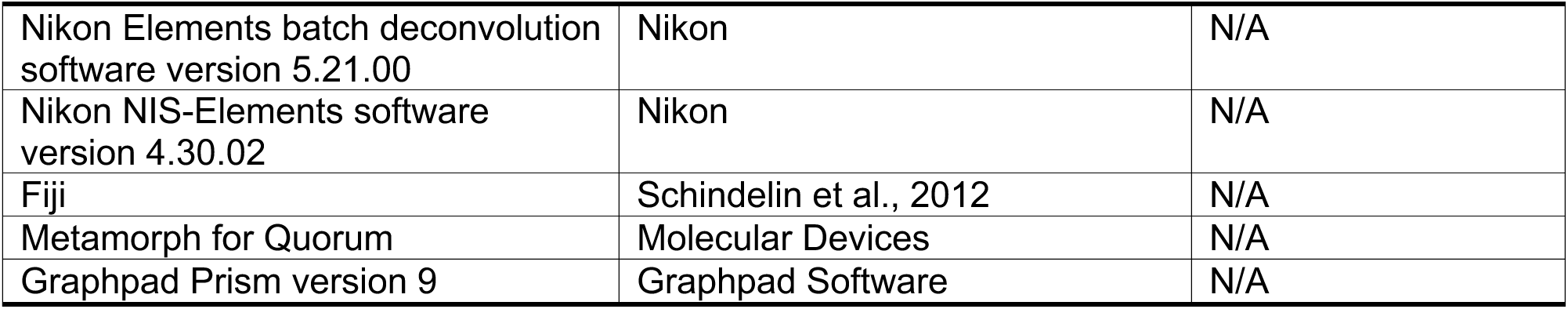

## Data availability

Requests for resources presented in this work should be submitted to DC.

## Acknowledgements

This work was supported by a grant from the National Institutes of Health #GM051542 to DC. The authors acknowledge the following Shared Resources facilities (Bio-Rad ChemiDoc MP) at the Dartmouth Cancer Center with NCI Cancer Center Support Grant 5P30 CA023108-40.

## Author information

SB and DC: conceptualization, writing original draft, review and editing, and validation. SB: investigation, methodology, formal analysis, and data curation. DC: funding acquisition, resources, project administration, and supervision.

## Ethics declarations

The authors declare that they have no conflicts of interest

## Figure Legends for Supplemental Data

**Figure S1:**
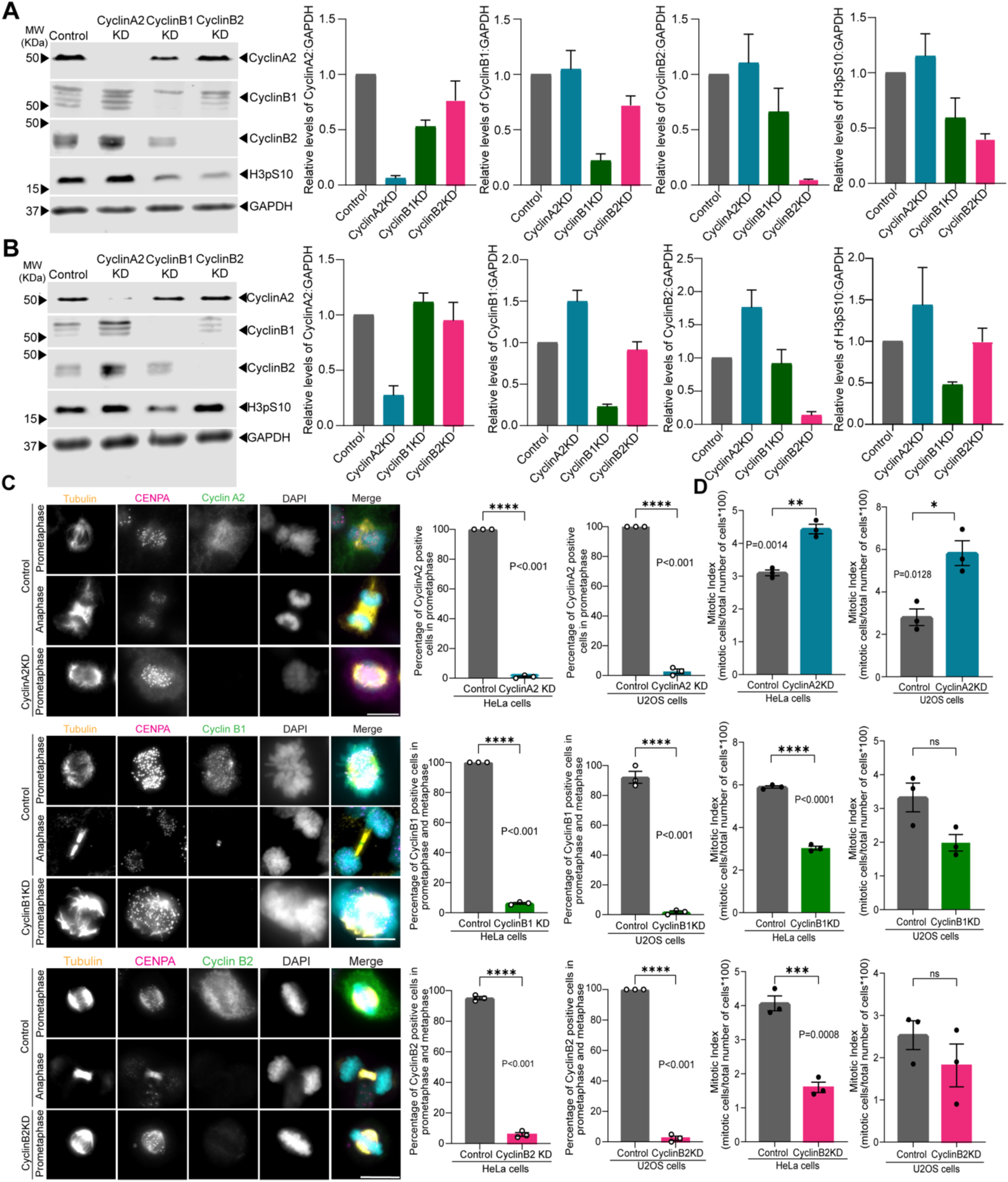
Validating Cyclin depletion in mitotic cells. A. Western blots from asynchronous HeLa cell populations for Cyclin A2, Cyclin B1, Cyclin B2, H3pS10 (mitotic marker) and GAPDH (loading control) for Control, Cyclin A2 knockdown (KD), Cyclin B1 KD and Cyclin B2 KD conditions. Quantification of the protein levels of Cyclin A2, Cyclin B1, Cyclin B2 and H3pS10 relative to GAPDH are shown in bar graphs from the blots. The relative protein levels were measured for each protein individually per condition represented by bars, where Control (grey bar), Cyclin A2 KD (blue bar), CyclinB1 KD (green bar) and Cyclin B2 KD (magenta) cells. (N=3) Error bars represent SEM. B. Western blots from asynchronous U2OS cell populations for Cyclin A2, Cyclin B1, Cyclin B2, H3pS10 (mitotic marker) and GAPDH (loading control) for Control, Cyclin A2 knockdown (KD), Cyclin B1 KD and Cyclin B2 KD conditions. Quantification of the protein levels of Cyclin A2, Cyclin B1, Cyclin B2 and H3pS10 relative to GAPDH are shown in bar graphs from the blots. The relative protein levels were measured for each protein individually per condition represented by bars, where Control (grey bar), Cyclin A2 KD (blue bar), CyclinB1 KD (green bar) and Cyclin B2 KD (magenta) cells. (N=3) Error bars represent SEM. C. Immunofluorescence images of individual mitotic cells during prometaphase and anaphase, as indicated, following depletion of Cyclin A2 KD, Cyclin B1 or Cyclin B2. Cells were stained for Tubulin (yellow), CENPA (magenta), Cyclin A2 or Cyclin B1 or Cyclin B2 (green) and DAPI (DNA; grey). Scale bar, 10μm. Graphs show the percentage of prometaphase cells scored positive for Cyclin A2 in Control (grey) and Cyclin A2 KD (blue) in HeLa (left) and U20S (right) cells, and percentage of prometaphase and metaphase cells scored positive for either Cyclin B1 or Cyclin B2 in Control (grey) and Cyclin B1 KD (green) or Cyclin B2 KD (magenta). (N=3, n=60 cells scored per condition). Values from each independent experiment is represented as circles on the bars. Error bars (black) are representative of Standard Error Mean (SEM) values. T-test was performed. Representative IF images of Control (prometaphase and anaphase) and Cyclin B1 KD (prometaphase) HeLa cells stained for Tubulin (yellow), CENPA (magenta), Cyclin B1 (green) and DAPI (grey). Scale bar: 10μm. D. Mitotic index calculated for HeLa (top) and U2OS (bottom) cells in Cyclin A2 KD, Cyclin B1 KD or Cyclin B2 KD and compared to the Control. Mitotic Index = (Number of mitotic cells/Total number of cells) * 100. Mitotic cells were counted in ten different field of views for each independent experiment. (N=3, n∼1500 cells for each condition, where n= total number of cells across three independent replicates). Values from each independent experiment has been represented as circles on the bars. Error bars (black) are representative of Standard Error Mean (SEM) values. Two tailed t-test was performed.

**Figure S2:**
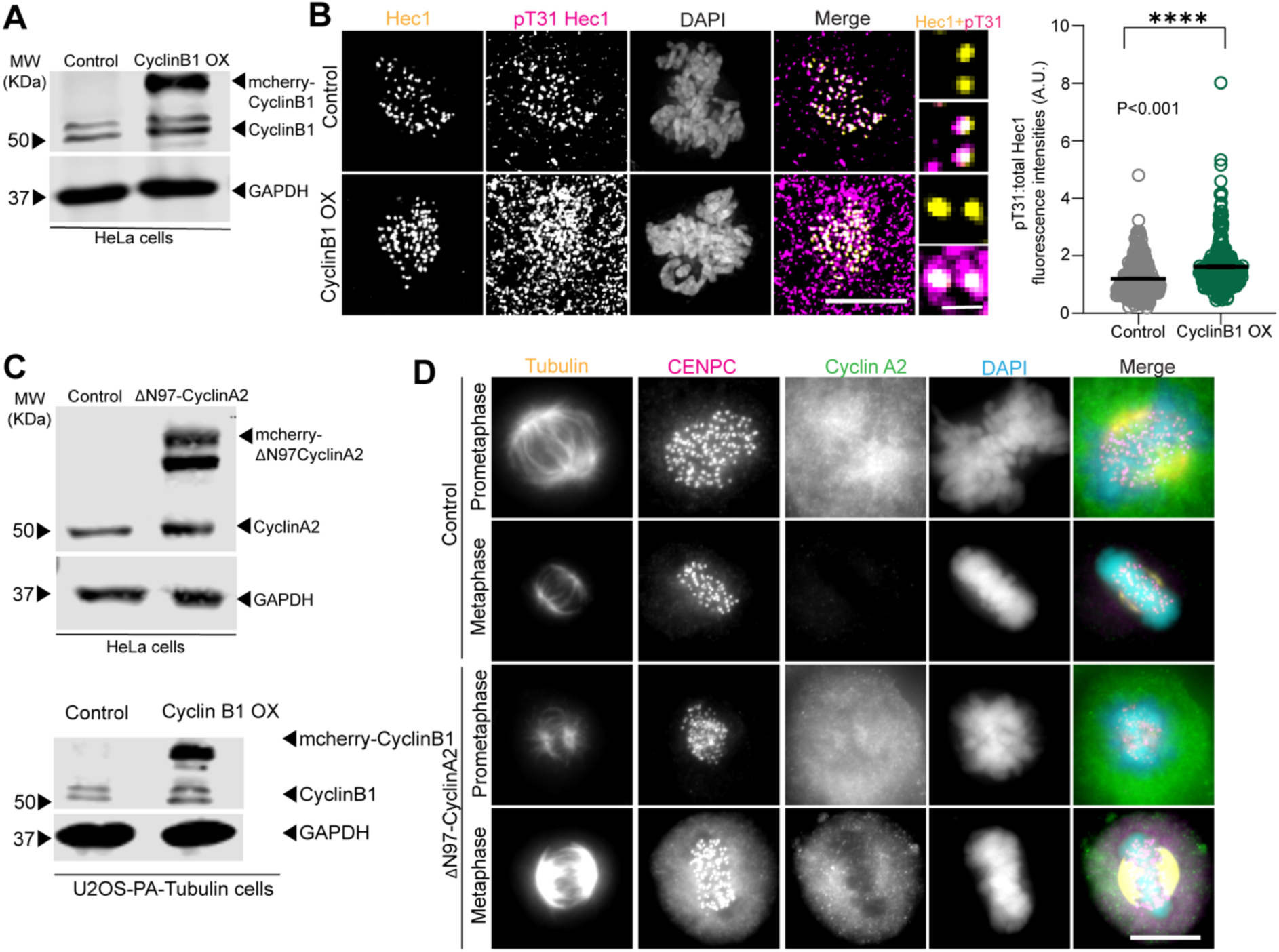
Validating Cyclin overexpression in mitotic cells. A. Western blots from asynchronous HeLa cell populations for Cyclin B1 and GAPDH (loading control) in Control and mCherry-tagged Cyclin B1 overexpressed (Cyclin B1 OX) cells. B. Immunofluorescence images of Control and CyclinB1 OX HeLa cells stained for Hec1(yellow), pT31 Hec1 (magenta) and DAPI (DNA; grey). Scale Bar, 10μm and 1μm for insets. Quantification of pT31 Hec1 kinetochore intensities relative to total Hec1 intensities in Control (grey) and Cyclin B1 OX (green) (N=3, n=300 kinetochores from 30 cells). Two tailed t-test was performed. C. Western blots from asynchronous HeLa cell populations for Cyclin A2 and GAPDH (loading control) are shown in Control and mCherry-tagged ΔN97 Cyclin A2 (ΔN97-Cyclin A2) expressing cells. Western blots from asynchronous U2OS cells expressing photoactivatable GFP-tagged alpha tubulin (U2OS-PH-tubulin) for mCherry-tagged Cyclin B1 and GAPDH (loading control). D. Immunofluorescence images of Control (prometaphase and metaphase) and ΔN97-Cyclin A2 (prometaphase and metaphase) expressing HeLa cells stained for Tubulin (yellow), CENPC (magenta), Cyclin A2 (green) and DAPI (DNA; grey). Scale Bar, 10μm.

**Figure S3:**
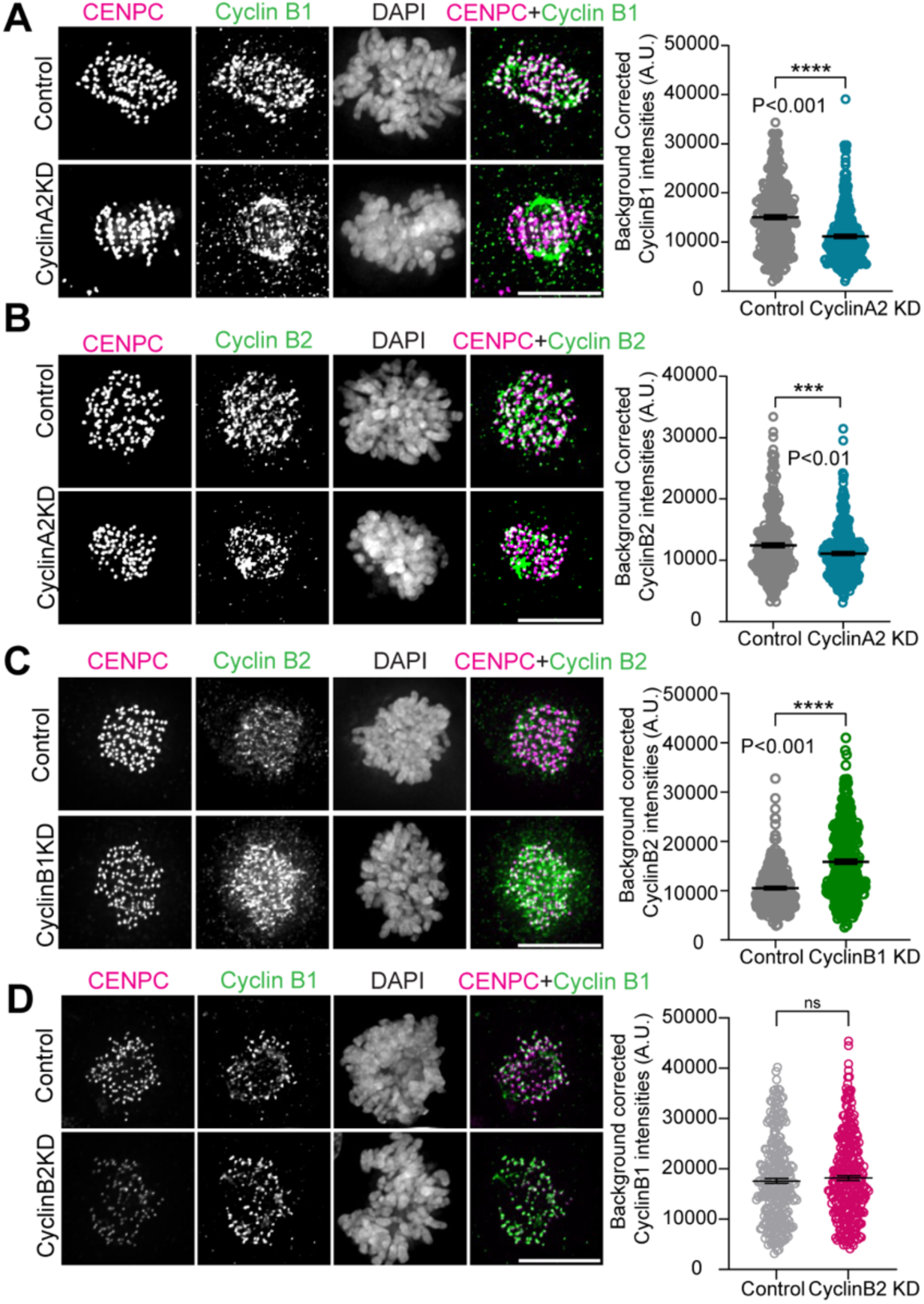
Cyclin isoform-specific regulation of cyclin localization in U2OS cells. A. Immunofluorescence images of Control and Cyclin A2 KD U2OS cells stained for CENPC, Cyclin B1 and DAPI (DNA). Quantification of Cyclin B1 intensities localized at the kinetochores in Control (grey) and Cyclin A2 KD (blue) cells (N=3, n=300 kinetochores from 30 cells). Two tailed t-test was performed. B. Immunofluorescence images of Control and Cyclin A2 U2OS cells stained for CENPC, Cyclin B2 and DAPI (DNA). Quantification of Cyclin B2 intensities localized at the kinetochores in Control (grey) and Cyclin A2 KD (blue) cells (N=3, n=300 kinetochores from 30 cells). Two tailed t-test was performed. C. Immunofluorescence images of Control and CyclinB1 KD U2OS cells stained for CENPC, Cyclin B2 and DAPI (DNA). Quantification of Cyclin B2 intensities localized at the kinetochores in Control (grey) and Cyclin B1 KD (green) cells (N=3, n=300 kinetochores from 30 cells). Two tailed t-test was performed. D. Immunofluorescence images of Control and CyclinB2 KD U2OS cells stained for CENPC, Cyclin B1 and DAPI (DNA). Quantification of Cyclin B1 intensities localized at the kinetochores in Control (grey) and Cyclin B2 KD (magenta) cells (N=3, n=300 kinetochores from 30 cells). Two tailed t-test was performed. Scale Bar, 10μm in all image panels.

**Figure S4:**
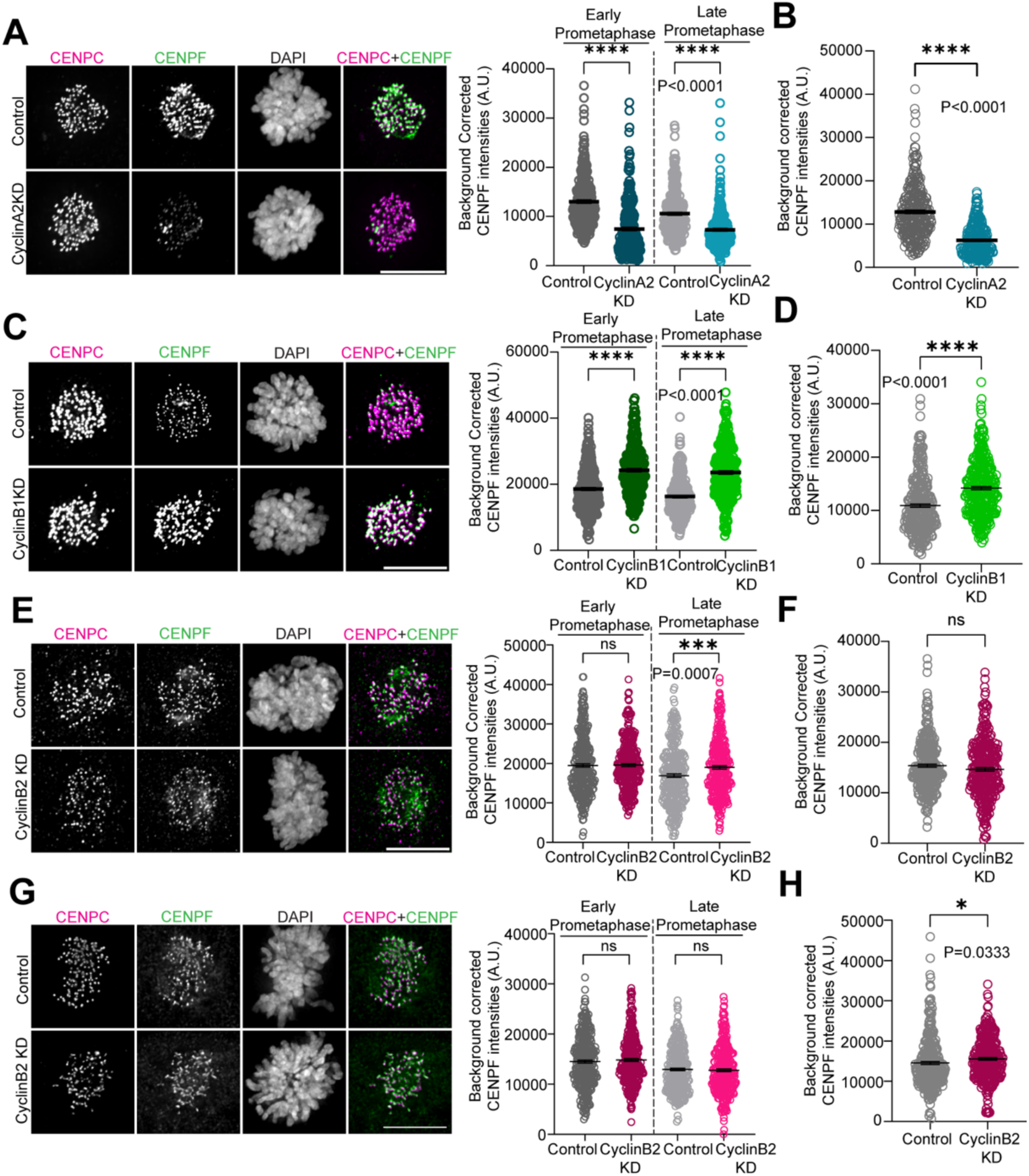
Cyclin isoform-specific localization of CENPF in U2OS cells. A. Immunofluorescence images of Control and Cyclin A2 KD cells during prometaphase stained for CENPC, CENPF and DAPI (DNA). Quantification of CENPF intensities localized at the kinetochores in control (grey) and Cyclin A2 KD (blue) cells in early and late prometaphase (N=3, n=300 kinetochores from 30 cells). One-way Anova was performed. B. Quantification of CENPF intensities localized at the kinetochores in control (grey) and Cyclin A2 KD (blue) cells (N=3, n=300 kinetochores from 30 cells). Cells were treated with 3.3μM nocodazole for 2 hours prior to fixation to deplete spindle microtubules. Student t-test was performed. C. Immunofluorescence images of Control and Cyclin B1 KD cells during prometaphase stained for CENPC, CENPF and DAPI (DNA). Quantification of CENPF intensities localized at the kinetochores in control (grey) and Cyclin B1 KD (green) cells in early and late prometaphase (N=3, n=300 kinetochores from 30 cells). One-way Anova was performed. D. Quantification of CENPF intensities localized at the kinetochores in control (grey) and Cyclin B1 KD (green) cells (N=3, n=300 kinetochores from 30 cells). Cells were treated with 3.3μM nocodazole for 2 hours prior to fixation to deplete spindle microtubules. Student t-test was performed. E. Immunofluorescence images of Control and Cyclin B2 KD HeLa cells during prometaphase stained for CENPC, CENPF and DAPI (DNA). Quantification of CENPF intensities localized at the kinetochores in control (grey) and Cyclin B2 KD (magenta) cells in early and late prometaphase (N=3, n=300 kinetochores from 30 cells). One-way Anova was performed. F. Quantification of CENPF intensities localized at the kinetochores in control (grey) and Cyclin B2 KD (magenta) cells (N=3, n=300 kinetochores from 30 cells). HeLa cells were treated with 3.3μM nocodazole for 2 hours prior to fixation to deplete spindle microtubules. Student t-test was performed. G. Immunofluorescence images of Control and Cyclin B2 KD U2OS cells during prometaphase stained for CENPC, CENPF and DAPI (DNA). Quantification of CENPF intensities localized at the kinetochores in control (grey) and Cyclin B2 KD (magenta) cells in early and late prometaphase (N=3, n=300 kinetochores from 30 cells). One-way Anova was performed. H. Quantification of CENPF intensities localized at the kinetochores in control (grey) and Cyclin B2 KD (magenta) cells (N=3, n=300 kinetochores from 30 cells). U2OS cells were treated with 3.3μM nocodazole for 2 hours prior to fixation to deplete spindle microtubules. Student t-test was performed. Scale bar, 10μm in all image panels

## References

Alfonso-Pérez T, Hayward D, Holder J, Gruneberg U & Barr FA (2019) MAD1-dependent recruitment of CDK1-CCNB1 to kinetochores promotes spindle checkpoint signaling. J Cell Biol 218: 1108–1117

Ali A & Stukenberg PT (2023) Aurora kinases: Generators of spatial control during mitosis. Front Cell Dev Biol 11

Allan LA, Camacho Reis M, Ciossani G, Huis In’t Veld PJ, Wohlgemuth S, Kops GJ, Musacchio A & Saurin AT (2020) Cyclin B1 scaffolds MAD1 at the kinetochore corona to activate the mitotic checkpoint. EMBO J 39: e103180

Al-Rawi A, Kaye E, Korolchuk S, Endicott JA & Ly T (2023) Cyclin A and Cks1 promote kinase consensus switching to non-proline-directed CDK1 phosphorylation. Cell Rep 42: 112139

Auckland P, Roscioli E, Coker HLE & McAinsh AD (2020) CENP-F stabilizes kinetochore-microtubule attachments and limits dynein stripping of corona cargoes. J Cell Biol 219: e201905018

Bentley AM, Normand G, Hoyt J & King RW (2007) Distinct sequence elements of cyclin B1 promote localization to chromatin, centrosomes, and kinetochores during mitosis. Mol Biol Cell 18: 4847–4858

Broad AJ & DeLuca JG (2020) The right place at the right time: Aurora B kinase localization to centromeres and kinetochores. Essays Biochem 64: 299–311

Brown NR, Lowe ED, Petri E, Skamnaki V, Antrobus R & Johnson LN (2007) Cyclin B and cyclin A confer different substrate recognition properties on CDK2. Cell Cycle Georget Tex 6: 1350–1359

Carmena M, Wheelock M, Funabiki H & Earnshaw WC (2012) The chromosomal passenger complex (CPC): from easy rider to the godfather of mitosis. Nat Rev Mol Cell Biol 13: 789–803

Cheeseman IM (2014) The kinetochore. Cold Spring Harb Perspect Biol 6: a015826

Chen Q, Zhang X, Jiang Q, Clarke PR & Zhang C (2008) Cyclin B1 is localized to unattached kinetochores and contributes to efficient microtubule attachment and proper chromosome alignment during mitosis. Cell Res 18: 268–280

Chen Y-C, Kilic E, Wang E, Rossman W & Suzuki A (2025) CENcyclopedia: dynamic landscape of kinetochore architecture throughout the cell cycle. Nat Commun 16: 7676

Chung HJ, Park JE, Lee NS, Kim H & Jang C-Y (2016) Phosphorylation of Astrin Regulates Its Kinetochore Function. J Biol Chem 291: 17579–17592

Cimini D, Wan X, Hirel CB & Salmon ED (2006) Aurora kinase promotes turnover of kinetochore microtubules to reduce chromosome segregation errors. Curr Biol CB 16: 1711–1718

Conti D, Gul P, Islam A, Martín-Durán JM, Pickersgill RW & Draviam VM (2019) Kinetochores attached to microtubule-ends are stabilised by Astrin bound PP1 to ensure proper chromosome segregation. eLife 8: e49325

Cordeiro MH, Smith RJ & Saurin AT (2018) A fine balancing act: A delicate kinase-phosphatase equilibrium that protects against chromosomal instability and cancer. Int J Biochem Cell Biol 96: 148–156

Crncec A, Lau HW, Ng LY, Ma HT, Mak JPY, Choi HF, Yeung TK & Poon RYC (2025) Plasticity of mitotic cyclins in promoting the G2-M transition. J Cell Biol 224: e202409219

Cui H, Loftus KM, Noell CR & Solmaz SR (2018) Identification of Cyclin-dependent Kinase 1 Specific Phosphorylation Sites by an In Vitro Kinase Assay. J Vis Exp: 57674

D’Angiolella V, Mari C, Nocera D, Rametti L & Grieco D (2003) The spindle checkpoint requires cyclin-dependent kinase activity. Genes Dev 17: 2520–2525

Dumitru AMG, Rusin SF, Clark AEM, Kettenbach AN & Compton DA (2017) Cyclin A/Cdk1 modulates Plk1 activity in prometaphase to regulate kinetochore-microtubule attachment stability. eLife 6: e29303

den Elzen N & Pines J (2001) Cyclin a Is Destroyed in Prometaphase and Can Delay Chromosome Alignment and Anaphase. J Cell Biol 153: 121–136

Ferrell JE (2002) Self-perpetuating states in signal transduction: positive feedback, double-negative feedback and bistability. Curr Opin Cell Biol 14: 140–148

Fisher DL & Nurse P (1996) A single fission yeast mitotic cyclin B p34cdc2 kinase promotes both S-phase and mitosis in the absence of G1 cyclins. EMBO J 15: 850–860

Friese A, Faesen AC, Huis in ‘t Veld PJ, Fischböck J, Prumbaum D, Petrovic A, Raunser S, Herzog F & Musacchio A (2016) Molecular requirements for the inter-subunit interaction and kinetochore recruitment of SKAP and Astrin. Nat Commun 7: 11407

Fung TK, Ma HT & Poon RYC (2007) Specialized roles of the two mitotic cyclins in somatic cells: cyclin A as an activator of M phase-promoting factor. Mol Biol Cell 18: 1861–1873

Fung TK & Poon RYC (2005) A roller coaster ride with the mitotic cyclins. Semin Cell Dev Biol 16: 335–342

Furuno N, den Elzen N & Pines J (1999) Human cyclin A is required for mitosis until mid prophase. J Cell Biol 147: 295–306

Geraghty Z, Barnard C, Uluocak P & Gruneberg U (2021) The association of Plk1 with the astrin–kinastrin complex promotes formation and maintenance of a metaphase plate. J Cell Sci 134: jcs251025

Godek KM, Kabeche L & Compton DA (2015) Regulation of kinetochore–microtubule attachments through homeostatic control during mitosis. Nat Rev Mol Cell Biol 16: 57–64

Gong D & Ferrell JE (2010a) The Roles of Cyclin A2, B1, and B2 in Early and Late Mitotic Events. Mol Biol Cell 21: 3149–3161

Gong D & Ferrell JE (2010b) The Roles of Cyclin A2, B1, and B2 in Early and Late Mitotic Events. Mol Biol Cell 21: 3149–3161

Gong D, Pomerening JR, Myers JW, Gustavsson C, Jones JT, Hahn AT, Meyer T & Ferrell JE (2007) Cyclin A2 Regulates Nuclear-Envelope Breakdown and the Nuclear Accumulation of Cyclin B1. Curr Biol 17: 85–91

Hayward D, Alfonso-Pérez T, Cundell MJ, Hopkins M, Holder J, Bancroft J, Hutter LH, Novak B, Barr FA & Gruneberg U (2019) CDK1-CCNB1 creates a spindle checkpoint–permissive state by enabling MPS1 kinetochore localization. J Cell Biol 218: 1182–1199

Hégarat N, Crncec A, Suarez Peredo Rodriguez MF, Echegaray Iturra F, Gu Y, Busby O, Lang PF, Barr AR, Bakal C, Kanemaki MT, et al (2020) Cyclin A triggers Mitosis either via the Greatwall kinase pathway or Cyclin B. EMBO J 39

Heinzle C, Höfler A, Yu J, Heid P, Kremer N, Schunk R, Stengel F, Bange T, Boland A & Mayer TU (2025) Positively charged specificity site in cyclin B1 is essential for mitotic fidelity. Nat Commun 16: 853

Kabeche L & Compton DA (2013) Cyclin A regulates kinetochore microtubules to promote faithful chromosome segregation. Nature 502: 110–113

Kalaszczynska I, Geng Y, Iino T, Mizuno S, Choi Y, Kondratiuk I, Silver DP, Wolgemuth DJ, Akashi K & Sicinski P (2009) Cyclin A Is Redundant in Fibroblasts but Essential in Hematopoietic and Embryonic Stem Cells. Cell 138: 352–365

Kern DM, Monda JK, Su K-C, Wilson-Kubalek EM & Cheeseman IM (2017) Astrin-SKAP complex reconstitution reveals its kinetochore interaction with microtubule-bound Ndc80. eLife 6: e26866

Kops GJPL & Gassmann R (2020) Crowning the Kinetochore: The Fibrous Corona in Chromosome Segregation. Trends Cell Biol 30: 653–667

Krenn V & Musacchio A (2015) The Aurora B Kinase in Chromosome Bi-Orientation and Spindle Checkpoint Signaling. Front Oncol 5: 225

Kucharski TJ, Hards R, Vandal SE, Abad MA, Jeyaprakash AA, Kaye E, Al-Rawi A, Ly T, Godek KM, Gerber SA, et al (2022) Small changes in phospho-occupancy at the kinetochore-microtubule interface drive mitotic fidelity. J Cell Biol 221: e202107107

Liu D & Lampson MA (2009) Regulation of kinetochore-microtubule attachments by Aurora B kinase. Biochem Soc Trans 37: 976–980

Liu S, Yuan X, Gui P, Liu R, Durojaye O, Hill DL, Fu C, Yao X, Dou Z & Liu X (2022) Mad2 promotes Cyclin B2 recruitment to the kinetochore for guiding accurate mitotic checkpoint. EMBO Rep 23: e54171

Lowe ED, Tews I, Cheng KY, Brown NR, Gul S, Noble MEM, Gamblin SJ & Johnson LN (2002) Specificity determinants of recruitment peptides bound to phospho-CDK2/cyclin A. Biochemistry 41: 15625–15634

Mack GJ & Compton DA (2001) Analysis of mitotic microtubule-associated proteins using mass spectrometry identifies astrin, a spindle-associated protein. Proc Natl Acad Sci U S A 98: 14434–14439

Magidson V, Paul R, Yang N, Ault JG, O’Connell CB, Tikhonenko I, McEwen BF, Mogilner A & Khodjakov A (2015) Adaptive changes in the kinetochore architecture facilitate proper spindle assembly. Nat Cell Biol 17: 1134–1144

Maller JL (1991) Mitotic control. Curr Opin Cell Biol 3: 269–275

Manning AL, Bakhoum SF, Maffini S, Correia-Melo C, Maiato H & Compton DA (2010) CLASP1, astrin and Kif2b form a molecular switch that regulates kinetochore-microtubule dynamics to promote mitotic progression and fidelity. EMBO J 29: 3531–3543

McAinsh AD & Kops GJPL (2023) Principles and dynamics of spindle assembly checkpoint signalling. Nat Rev Mol Cell Biol 24: 543–559

Murray AW (1993) Cell-cycle control: turning on mitosis. Curr Biol CB 3: 291–293

Murray AW (2004) Recycling the cell cycle: cyclins revisited. Cell 116: 221–234

Musacchio A & Salmon ED (2007) The spindle-assembly checkpoint in space and time. Nat Rev Mol Cell Biol 8: 379–393

Nurse P (2002) Cyclin dependent kinases and cell cycle control (nobel lecture). Chembiochem Eur J Chem Biol 3: 596–603

Pagliuca FW, Collins MO, Lichawska A, Zegerman P, Choudhary JS & Pines J (2011) Quantitative proteomics reveals the basis for the biochemical specificity of the cell-cycle machinery. Mol Cell 43: 406–417

Pellarin I, Dall’Acqua A, Favero A, Segatto I, Rossi V, Crestan N, Karimbayli J, Belletti B & Baldassarre G (2025) Cyclin-dependent protein kinases and cell cycle regulation in biology and disease. Signal Transduct Target Ther 10: 11

Pines J (1995) Cyclins and cyclin-dependent kinases: a biochemical view. Biochem J 308 ( Pt 3): 697–711

Rosas-Salvans M, Sutanto R, Suresh P & Dumont S (2022) The Astrin-SKAP Complex Reduces Friction at the Kinetochore-Microtubule Interface. Curr Biol CB 32: 2621–2631.e3

Roy B, Han SJY, Fontan AN, Jema S & Joglekar AP (2022) Aurora B phosphorylates Bub1 to promote spindle assembly checkpoint signaling. Curr Biol 32: 237–247.e6

Sacristan C, Ahmad MUD, Keller J, Fermie J, Groenewold V, Tromer E, Fish A, Melero R, Carazo JM, Klumperman J, et al (2018) Dynamic kinetochore size regulation promotes microtubule capture and chromosome biorientation in mitosis. Nat Cell Biol 20: 800–810

Saurin AT, van der Waal MS, Medema RH, Lens SMA & Kops GJPL (2011) Aurora B potentiates Mps1 activation to ensure rapid checkpoint establishment at the onset of mitosis. Nat Commun 2: 316

Schmidt A-K, Pudelko K, Boekenkamp J-E, Berger K, Kschischo M & Bastians H (2021) The p53/p73 - p21CIP1 tumor suppressor axis guards against chromosomal instability by restraining CDK1 in human cancer cells. Oncogene 40: 436–451

Schmidt JC, Kiyomitsu T, Hori T, Backer CB, Fukagawa T & Cheeseman IM (2010) Aurora B kinase controls the targeting of the Astrin-SKAP complex to bioriented kinetochores. J Cell Biol 191: 269–280

Scholey JM, Brust-Mascher I & Mogilner A (2003) Cell division. Nature 422: 746–752

Schulman BA, Lindstrom DL & Harlow E (1998) Substrate recruitment to cyclin-dependent kinase 2 by a multipurpose docking site on cyclin A. Proc Natl Acad Sci U S A 95: 10453–10458

Serpico AF & Grieco D (2020) Recent advances in understanding the role of Cdk1 in the Spindle Assembly Checkpoint. F1000Research 9: F1000 Faculty Rev-57

Shrestha RL, Conti D, Tamura N, Braun D, Ramalingam RA, Cieslinski K, Ries J & Draviam VM (2017) Aurora-B kinase pathway controls the lateral to end-on conversion of kinetochore-microtubule attachments in human cells. Nat Commun 8: 150

Silva Cascales H, Burdova K, Middleton A, Kuzin V, Müllers E, Stoy H, Baranello L, Macurek L & Lindqvist A (2021) Cyclin A2 localises in the cytoplasm at the S/G2 transition to activate PLK1. Life Sci Alliance 4: e202000980

Song X, Conti D, Shrestha RL, Braun D & Draviam VM (2021) Counteraction between Astrin-PP1 and Cyclin-B-CDK1 pathways protects chromosome-microtubule attachments independent of biorientation. Nat Commun 12: 7010

Tsukahara T, Tanno Y & Watanabe Y (2010) Phosphorylation of the CPC by Cdk1 promotes chromosome bi-orientation. Nature 467: 719–723

Valles SY, Bural S, Godek KM & Compton DA (2024) Cyclin A/Cdk1 promotes chromosome alignment and timely mitotic progression. Mol Biol Cell 35: ar141

Wolgemuth DJ, Lele KM, Jobanputra V & Salazar G (2004) The A-type cyclins and the meiotic cell cycle in mammalian male germ cells. Int J Androl 27: 192–199

Wu J, Larreategui-Aparicio A, Lambers MLA, Bodor DL, Klaasen SJ, Tollenaar E, de Ruijter-Villani M & Kops GJPL (2023) Microtubule nucleation from the fibrous corona by LIC1-pericentrin promotes chromosome congression. Curr Biol CB 33: 912–925.e6

Zhai Y, Kronebusch PJ & Borisy GG (1995) Kinetochore microtubule dynamics and the metaphase-anaphase transition. J Cell Biol 131: 721–734

Zhang Q-H, Yuen WS, Adhikari D, Flegg JA, FitzHarris G, Conti M, Sicinski P, Nabti I, Marangos P & Carroll J (2017) Cyclin A2 modulates kinetochore–microtubule attachment in meiosis II. J Cell Biol 216: 3133–3143

